# Autologous human immunocompetent white adipose tissue-on-chip

**DOI:** 10.1101/2021.08.08.455559

**Authors:** Julia Rogal, Raylin Xu, Julia Roosz, Claudia Teufel, Madalena Cipriano, Wiebke Eisler, Martin Weiss, Katja Schenke-Layland, Peter Loskill

## Abstract

Obesity and associated diseases, such as diabetes, have reached epidemic proportions globally. In the era of ‘diabesity’ and due to its central role for metabolic and endocrine processes, adipose tissue (specifically white adipose tissue; WAT) has become a target of high interest for therapeutic strategies. To gain insights in cellular and molecular mechanisms of adipose (patho-)physiology, researchers traditionally relied on animal models since *in vitro* studies on human WAT are challenging due to the large size, buoyancy, and fragility of mature white adipocytes. Leveraging the Organ-on-Chip technology, we introduce a next-generation microphysiological *in vitro* model of human WAT based on a tailored microfluidic platform featuring vasculature-like perfusion. The platform integrates a 3D tissue comprising all major WAT-associated cellular components in an autologous manner, including not only mature adipocytes but also organotypic endothelial barriers and stromovascular cells featuring tissue-resident innate immune cells, specifically adipose tissue macrophages. This microphysiological tissue model recapitulates pivotal WAT functions, such as energy storage and mobilization as well as endocrine and immunomodulatory activities. The combination of all individual cell types with extra cellular matrix-like hydrogels in a precisely controllable bottom-up approach enables the generation of a multitude of replicates from the same donors circumventing issues of inter-donor variability and paving the way for personalized medicine. Moreover, it allows to adjust the model’s degree of complexity to fit a specific purpose via a flexible mix-and-match approach with different cell component modules. This novel WAT-on-chip system constitutes a human-based, autologous and immunocompetent *in vitro* model of adipose tissue that recapitulates almost full tissue heterogeneity. In the future, the new WAT-on-chip model can become a powerful tool for human-relevant research in the field of metabolism and its associated diseases as well as for compound testing and personalized- and precision medicine applications.

## Introduction

Obesity, defined by a body mass index (BMI) of 30 or above, has reached epidemic proportions globally. About 13% of the world’s adult population was obese in 2016 (World Health Organization, 2021) – and this number has continued to rise. Marked by a state of low-grade chronic inflammation, obesity is a well-recognized risk factor for a myriad of co-morbidities, amongst them type 2 diabetes mellitus (T2DM), cardiovascular and neurodegenerative diseases, at least 13 different types of cancer (National Cancer Institute at the National Institutes of Health, 2020), and infectious diseases [e.g. COVID19 (Andrade et al., 2021; Hornung et al., 2021)]. Moreover, being obese directly impacts the immune system’s ability to respond to infections (Alarcon et al., 2021; Hornung et al., 2021). Therefore, in the era of ‘diabesity’ and due to its central role for metabolic and endocrine processes, adipose tissue has become a target of high interest for therapeutic strategies against various diseases.

Adipose tissues can be categorized into white adipose tissue (WAT), brown adipose tissue (BAT), brite/beige adipose tissue, and pink adipose tissue. Each tissue type is morphologically distinct and performs unique functions (Corrêa et al., 2019). In this study, we focus on WAT. White adipocytes are integral components of WAT and highly specialized to lipid metabolism. Unlike any other cell type, they can take up and store vast amounts of lipids without being damaged. Moreover, white adipocytes are well equipped to sense and govern the body’s energy status. These cells make up about 90% of WAT volume but less than 50% of cellular content (Corvera, 2021). The remaining WAT-associated cell populations are broadly pooled as stromal vascular fraction (SVF). These stromovascular cells include adipose-derived mesenchymal stem cells (AdMSCs), adipocyte and vascular progenitors, fibroblasts, as well as tissue-resident immune cells. Crosstalk between stromovascular cells and adipocytes considerably contributes to modulation of immune responses (Morigny et al., 2021; Sun et al., 2011). Dysfunction of both storage and endocrine WAT activity can have systemic consequences. The close connection between WAT and the immune system comes as no surprise. The most frequent immune cell populations in WAT are adipose tissue macrophages (ATMs), eosinophils, innate lymphoid cells, T cells, and B cells (Eberl et al., 2015; Guzik et al., 2017; Han et al., 2017; Patel et al., 2020; Srikakulapu and McNamara, 2020). Typically, adipose tissue immune cells control integrity and hormone sensitivity of adipocytes (Reilly and Saltiel, 2017). Yet, in response to overnutrition, adipocytes expand in number (hyperplasia) and size (hypertrophy) and eventually unleash a cascade of inflammatory events. Alongside adipocyte-associated functional changes, such as disturbed fatty acid (FA) metabolism or increased insulin resistance, this adipose tissue inflammation is marked by an accelerated immune cell infiltration. For instance, ATMs constitute about 5%-10% of the SVF in healthy humans but up to 50% in obesity (Russo and Lumeng, 2018; Weisberg et al., 2003). Consequently, WAT has become highly relevant for studies on systemic immunometabolism (Lercher et al., 2020).

Gaining human-relevant cellular and molecular insights in adipose (patho-)physiology, however, has traditionally been limited by several aspects: (i) *in vivo* human studies on mechanistic pathways usually entail unacceptable health risks. Thus, a large part of our understanding regarding human WAT function builds on clinical, mostly systemic, observations and genome-wide association studies (GWAS). (ii) Even though *in vivo* animal models allow for more flexibility regarding depth of biological level and degree of experimental interventions, their predictive value for humans is limited. There are major discrepancies between mice and humans, especially when it comes to metabolism and immunology (Greek and Menache, 2013; Mestas and Hughes, 2004; Reitman, 2018; Remick, 2005; van der Worp et al., 2010). (iii) *In vitro* studies on human WAT can be challenging due to the large size, buoyancy, and fragility of mature white adipocytes; rendering conventional cell culture methods unsuitable. Additionally, studies using WAT explants frequently encounter difficulties caused by hypoxia or inflammation (Fain et al., 2010; Gesta et al., 2003). Thus, many adipose *in vitro* studies utilized *in vitro* differentiation of adipocyte progenitors. However, so far, the maturity of these differentiated adipocytes does not adequately reflect the biology and functionality of mature adipocytes (Bahmad et al., 2020; Li and Easley, 2018; Volz et al., 2019).

As a consequence, compared to other organ systems, research on *in vitro* adipose tissue models has been rather sparse. Additionally, the predominant intention behind adipose tissue engineering has been the development of large-scale tissue grafts for regenerative medicine, rather than studies on adipose (patho-)physiological mechanisms. Still, several efforts have been made to come up with advanced long-term tissue culture models that can circumvent the restraints in mature adipocyte handling and culturability. 3D biomaterial scaffolds are often utilized to provide protection and a certain degree of structural stability (Abbott et al., 2018, 2016b, 2016a; Huber et al., 2016; Louis et al., 2019). Along the same line, structurally supported *in vitro* cultures have been achieved via sandwiching strategies, trapping mature adipocytes between SVF cell sheets (Lau et al., 2018), and sophisticated versions of ceiling cultures taking advantage of adipocyte buoyancy (Harms et al., 2019). While these approaches considerably contributed to the longevity of mature adipocytes *in vitro*, they still fall short on recapitulating key aspects of the adipose tissue microenvironment including vascular perfusion, cell-cell interactions as well as immune components.

In recent years, the Organ-on-Chip (OoC) technology has become a powerful tool for building *in vitro* culture systems that are reflective of human physiology. Combining microfabrication techniques and tissue engineering, OoCs emulate *in vivo* functionality of a certain organ or tissue at the smallest possible scale in a microfluidic platform. Alongside organ-specific 3D microenvironments, physiological cell-cell and cell-matrix interactions, one of the key features of OoCs is the vasculature-like perfusion; an aspect that is especially important for WAT *in vitro* culture, in view of its high metabolic and endocrine activity. Nevertheless, the current landscape of WAT-on-chip models is still scarce and shaped by *in vitro* differentiated adipocytes (Bahmad et al., 2020; Li and Easley, 2018; McCarthy et al., 2020). Despite some efforts to reflect insulin resistance or WAT immunoregulatory function, almost all WAT-on-chip models turn to differentiating AdMSCs/pre-adipocytes (Kongsuphol et al., 2019; Liu et al., 2019; Yang et al., 2021, 2020) or even murine preadipocytes as fundamental cellular components (Loskill et al., 2017; Tanataweethum et al., 2021, 2018; Zhu et al., 2018). Notably, there is a variety of microanalytical fluidic systems, which aim to interrogate adipocyte functionality using microfluidics approaches (Godwin et al., 2015; Hu et al., 2020; Li et al., 2018). While these analytical platforms integrate mature adipocytes and are powerful means to assess highly time-resolved adipocyte secretions, they are less suited for long-term culture of adipose tissue. To our best knowledge, the only OoC system, which is based on mature human adipocytes and adapted for long-term culture, is our previously published adipocyte-on-chip model (Rogal et al., 2020). Yet, this model integrates only adipocytes and thereby falls short on reflecting WAT’s full heterogeneity and consequent endocrine activities.

Here, we introduce a next-generation human WAT-on chip platform, which integrates all major WAT-associated cellular components in an autologous manner (figure 1). Mature adipocytes, together with stromovascular cells, or tissue-resident immune cells extracted from SVF, were encapsulated in a hydrogel matrix and injected into the microfluidic device’s tissue chambers. Media-perfused channels supplying the tissue chambers via diffusive exchange across a porous membrane were lined with tight layers of endothelium and served as travelling route for circulating immune cells (figure 1a). Besides the holistic reflection of the cellular composition of WAT, most importantly its immunocompetency, a key feature of our system is its fully autologous character (figure 1b). From skin biopsies with subcutaneous fat, we isolated mature adipocytes, SVF as well as microvascular endothelial cells (mvECs). In a further step, CD14^+^-cells, i.e., monocytes and macrophages, were separated from the SVF using magnetic activated cell sorting (MACS). For experiments on immune cell infiltration, T cells and CD14^+^-cells were derived from peripheral blood mononuclear cells (PBMCs), which were isolated from the biopsy donors’ blood. The individual cell types enabled us to build up a WAT model via a precisely controllable bottom-up approach that recapitulates pivotal WAT functions, such as energy storage and mobilization as well as endocrine and immunomodulatory activities. To adjust the model’s degree of complexity to fit a specific purpose, we introduce a flexible mix-and-match WAT-on-chip with different cell component modules.

**Figure 1.**
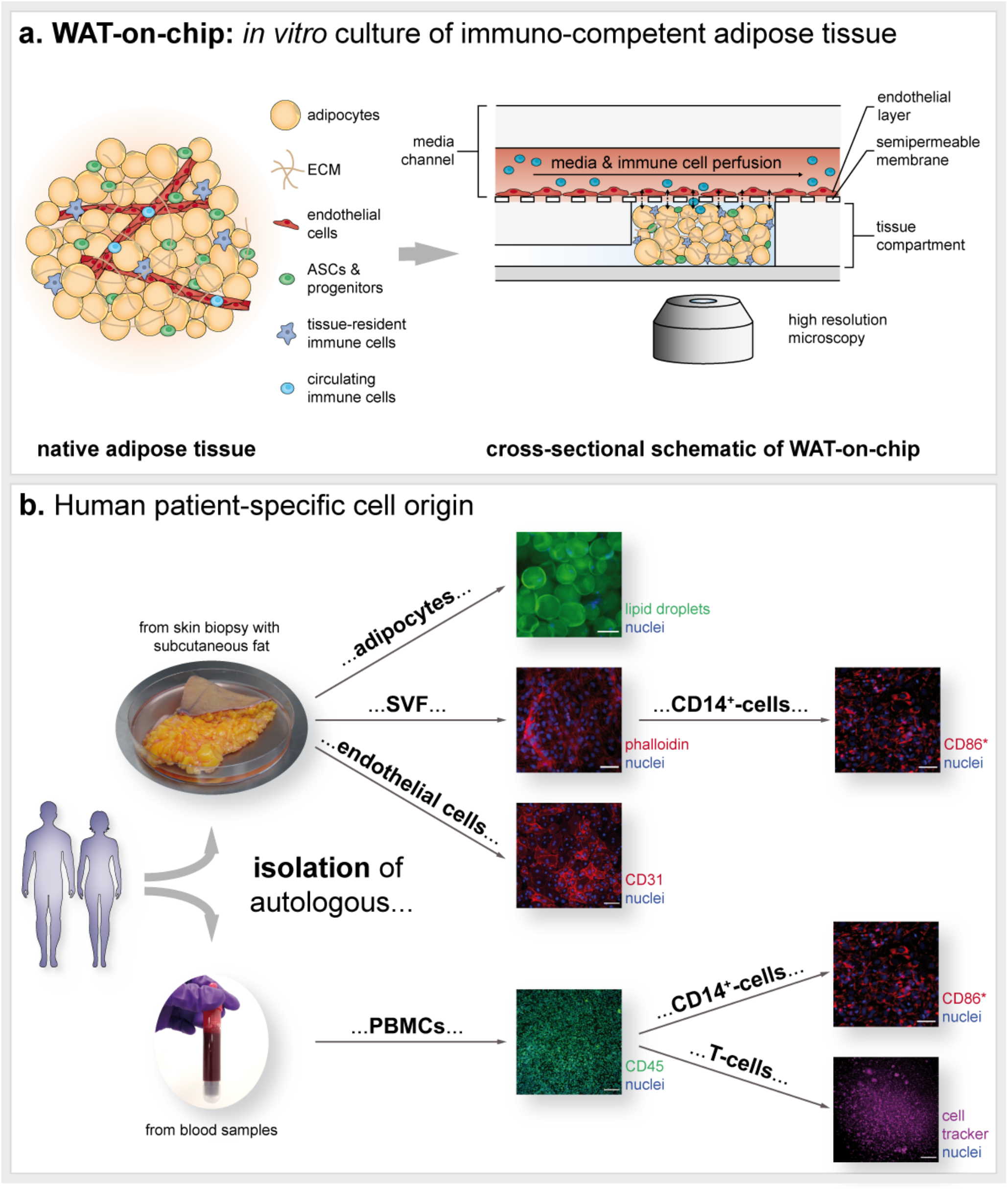
Concept of the human patient-specific WAT-on-chip model. (a) Schematic of WAT *in vivo* anatomy and integration of all cellular components into the microfluidic platform: Mature adipocytes, progenitors, stem cells, and tissue-resident immune cells are encapsulated in a hydrogel and cultured in the chips’ tissue compartment. Microvascular endothelial cells (mvECs) are seeded via the media channel onto the membrane shielding tissue chambers from the constant perfusion. To study immune cell recruitment, circulating immune cells were perfused through the media channels. (b) Patient-specific cell sources for building the WAT-on-chip model: Mature human adipocytes, mvECs as well as cells from the stromal vascular fraction (SVF), including tissue-resident immune cells such as CD14^+^-cells, are isolated from skin biopsies with subcutaneous fat. Circulating immune cells, such as T-cells, are retrieved by isolating PBMCs from the patients’ blood. Scale bars equal 100 µm (adipocytes, CD31, CD45 and T cell visualization) or 50 µm (SVF and CD14^+^-cell visualization). *CD14+-cells are isolated using magnetic activated cell sorting (MACS) and could therefore not be stained for CD14 but for CD86, another marker expressed on macrophages.

## Results and Discussion

### Microfluidic platform specifically tailored to accommodate adipose tissue

The microfluidic platform used in this study is a customized system specifically tailored to the integration of WAT (cf. figures 1a and 2a). The device was fabricated from two microstructured polydimethylsiloxane (PDMS) layers that are separated by a semipermeable, porous polyethylene terephthalate (PET) membrane. The lower PDMS layer was patterned with channel- and chamber microstructures to form the tissue compartment. It is comprised of eight individual tissue chambers branching off a common injection channel at a 45° angle and a thin, high-resistance channel towards the outlet port of the tissue compartment. Via the micropores in the PET membrane, the tissue chambers are connected to a constant media perfusion through media channels molded into the upper PDMS layer (media compartment). To the other side, the tissue chambers are encased by glass coverslips to enable optimal visual accessibility of on-chip tissues. The tissue chambers are 1 mm in diameter and feature a height of 0.2 mm each, resulting in a total tissue volume of 1.26 µl for the eight tissue chambers (for detailed dimensions cf. table 1, Materials and Methods section). In addition to the cylindrical shape of the tissue chambers, all edges in the tissue compartment were rounded to avoid cell damage. The media perfusion was realized by a parallel arrangement of media channels bifurcating from a common media inlet port that later merge to meet in a common media outlet port. We chose a parallel media perfusion over a serial media perfusion to avoid crosstalk among the individual chambers.

**Figure 2.**
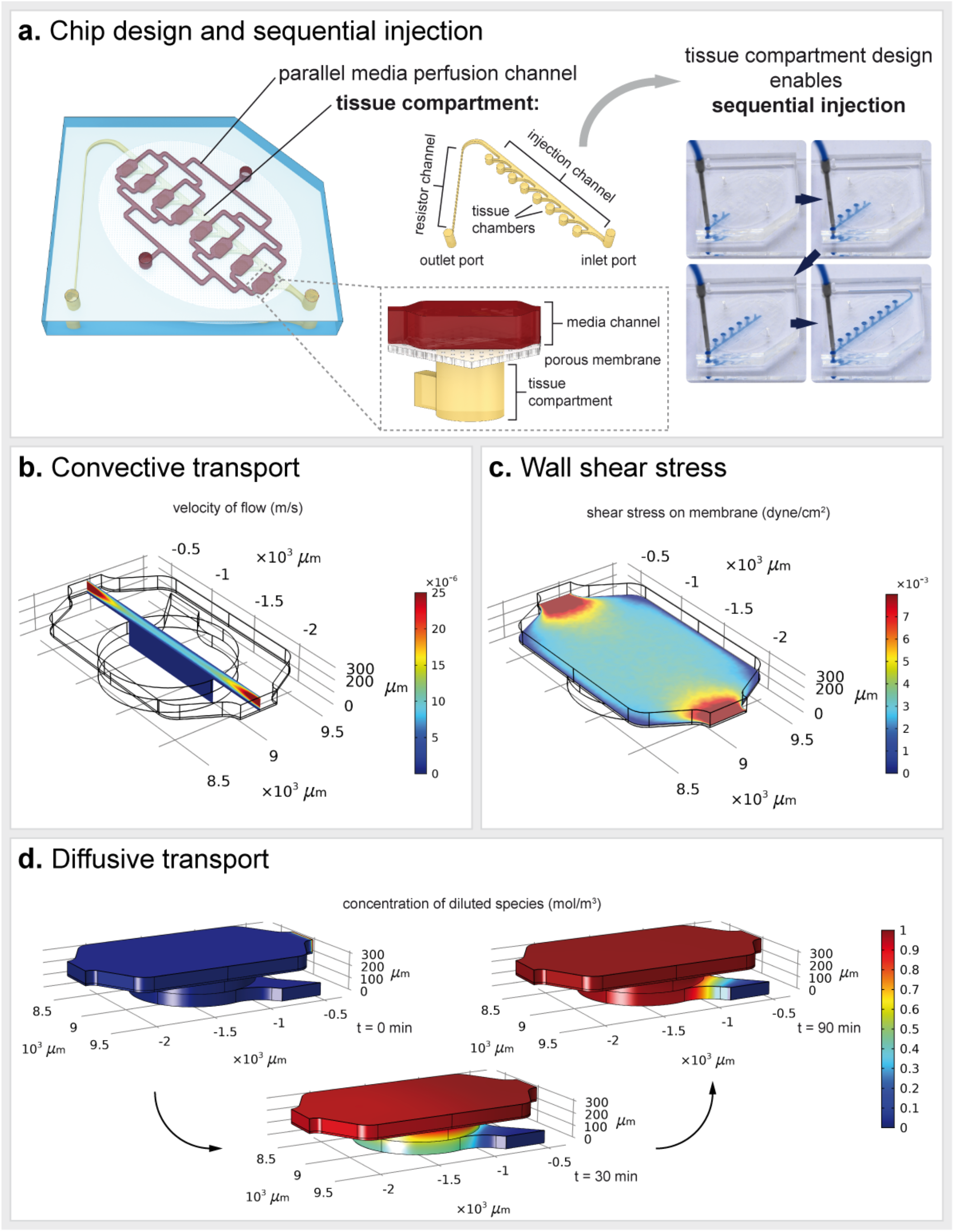
Characterization of the microfluidic platform. (a) Key characteristics of the WAT-on-chip platforms are (i) a parallel media perfusion channel to ensure equal media supply for each tissue chamber and (ii) a tissue compartment with eight separate tissue chambers. The design of the tissue compartment enables sequential injection of the tissue chambers. Computational fluid dynamic modeling revealed (b) a convective flow confined to the media channel, (c) low shear forces (∼ 0.002 – 0.006 dyn/cm^2^) on the membrane and (d) ensured diffusion of diluted species from the media flow into the tissue chambers. For all simulations, the flow rate was set to 2.5 µL/h per parallel channel, which results in 20 µL/h total flow rate, and the tissue compartment was assumed to be filled with a hydrogel.

**Table 1:** Dimensions of microfluidic WAT-on-chip platform.

| Tissue compartment |  |  |
| --- | --- | --- |
| Entire compartment | Area [$\text{mm}^2$] | 21.59 |
| | Volume [$\mu\text{l}$] | 2.80 |
| Dimensions of one chamber | Radius [mm] | 0.50 |
| | Area [$\text{mm}^2$] | 0.79 |
|  | Height [mm] | 0.20 |
| | Volume [$\mu\text{l}$] | 0.16 |
| Sum of eight chambers | Area [$\text{mm}^2$] | 6.28 |
| | Volume [$\mu\text{l}$] | 1.26 |
| Media compartment |  |  |
| Entire compartment | Area [$\text{mm}^2$] | 39.81 |
|  | Height [mm] | 0.10 |
| | Volume [$\mu\text{l}$] | 3.98 |

Besides housing the media channel compartment, the upper PDMS layer contains ports for tissue loading as well as media in- and outlet ports. Through the connection to an external syringe pump, we were able to precisely control the convective transport of substances, i.e., nutrients or drugs/compounds, to the tissue chambers as well as removal of metabolites or waste products from the on-chip tissues.

A key design feature of the platform is the architecture of the tissue compartment system, which enables a sequential injection procedure (figure 2a). Upon injecting the cell suspension through the tissue compartment’s inlet port, the chambers fill one after another following the path of lowest resistance (figure 2a); provided that the ports of the media compartment above are open. These injection properties are particularly favorable for handling human mature adipocytes: Owing to their large size and high lipid contents, these cells are extremely fragile. The sequential loading process prevents “overloading” of tissue chambers. Thereby, it protects the adipocytes from high pressures and potential damage during injection. Moreover, the technique facilitates a uniform loading and equal filling states among the chambers. The outlet of the tissue compartment is not only essential to the sequential injection principle; it also enables a clearing of the injection channel from surplus cells that did not fit into the tissue chambers anymore. For these remnant cells, a sufficient media supply could not be guaranteed, and cell death signals secreted by these remnant cells could negatively impact the perfused cells in the tissue chambers.

The separation of tissue chambers from the constant flow in the media channels by the porous membranes shields the tissue compartment from shear forces, as confirmed by computational fluid dynamics (CFD) modelling (figure 2b). The wall shear stress (WSS) on top of the membrane, above the hydrogel-filled tissue chambers, ranges between 3×10^-3^-4×10^-3^ dyn/cm^2^ (figure 2c). Yet, despite the membrane’s warding the tissue chambers from shear stress, sufficient nutrients reach the entire tissue chamber through diffusive transport across the membrane (figure 2c).

### Characterization of mature adipocytes-on-chip

After a general characterization of the microfluidic platform per se, we sought to investigate its suitability for the integration of human mature adipocytes suspended in a hydrogel matrix.

Human mature adipocytes were isolated from skin biopsies with subcutaneous adipose tissue and cultured overnight in flask-format. Prior to injection, the adipocytes were suspended in a hydrogel matrix and then injected into the tissue compartment. The hydrogel added a protective surrounding during injection and prevented buoyant adipocytes from floating to the top of the tissue chambers. Importantly, the integration of an adipocyte-surrounding matrix is physiologically relevant: *in vivo*, alterations in adipose tissue extracellular matrix (ECM) can lead to metabolic changes. An excess deposition of ECM, as is the case in obesity, was found to lead to an aggravation of insulin sensitivity (Lin et al., 2016), for instance. In our model, a synthetic hydrogel was used to achieve higher control and reproducibility compared to natural alternatives. Since collagens comprise the main ECM component in adipose tissue (Ruiz-Ojeda et al., 2019), we chose the HyStem^®^-C hydrogel, which is rich in denatured collagens providing cell attachment sites. Moreover, as a recent study reported that adipocytes in stiffer 3D matrices had increased pro-fibrotic gene expression profiles (Di Caprio and Bellas, 2020), we decreased the stiffness of the resulting hydrogel matrix by tailoring ratios of its components.

We characterized the adipocytes’ viability, morphology and functionality on-chip at different time points. Furthermore, we studied their response to ß-adrenergic-as well as pro-inflammatory stimulation. To assess impact of donor-variability, we also compared how cells from different donors (table 2, Materials and Methods) perform in the same experimental set-up.

**Table 2.**
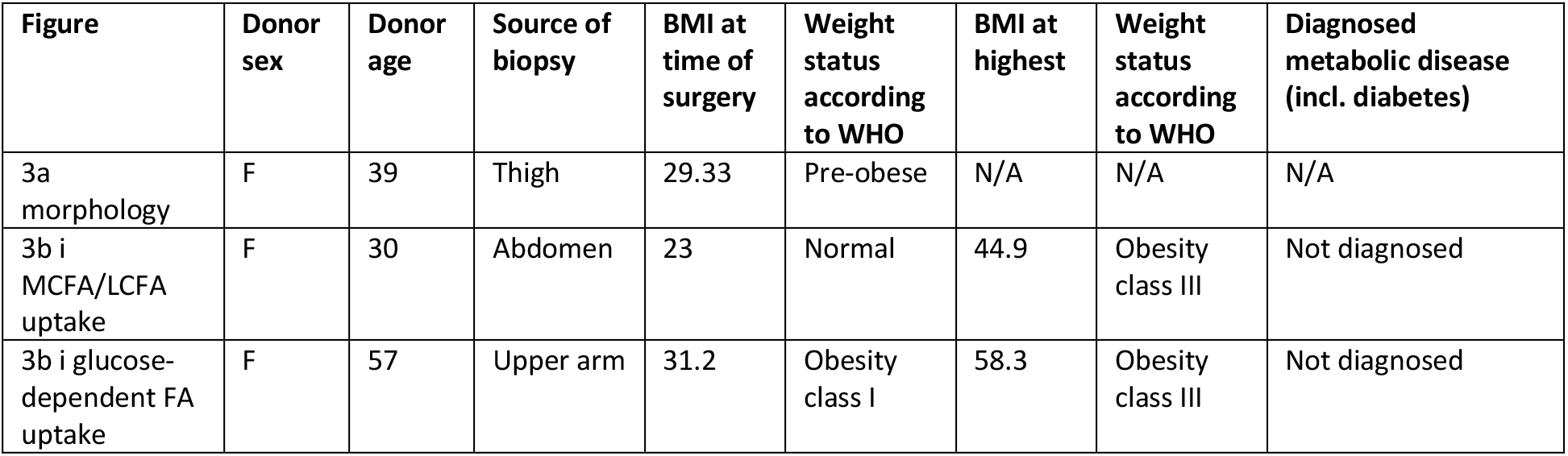
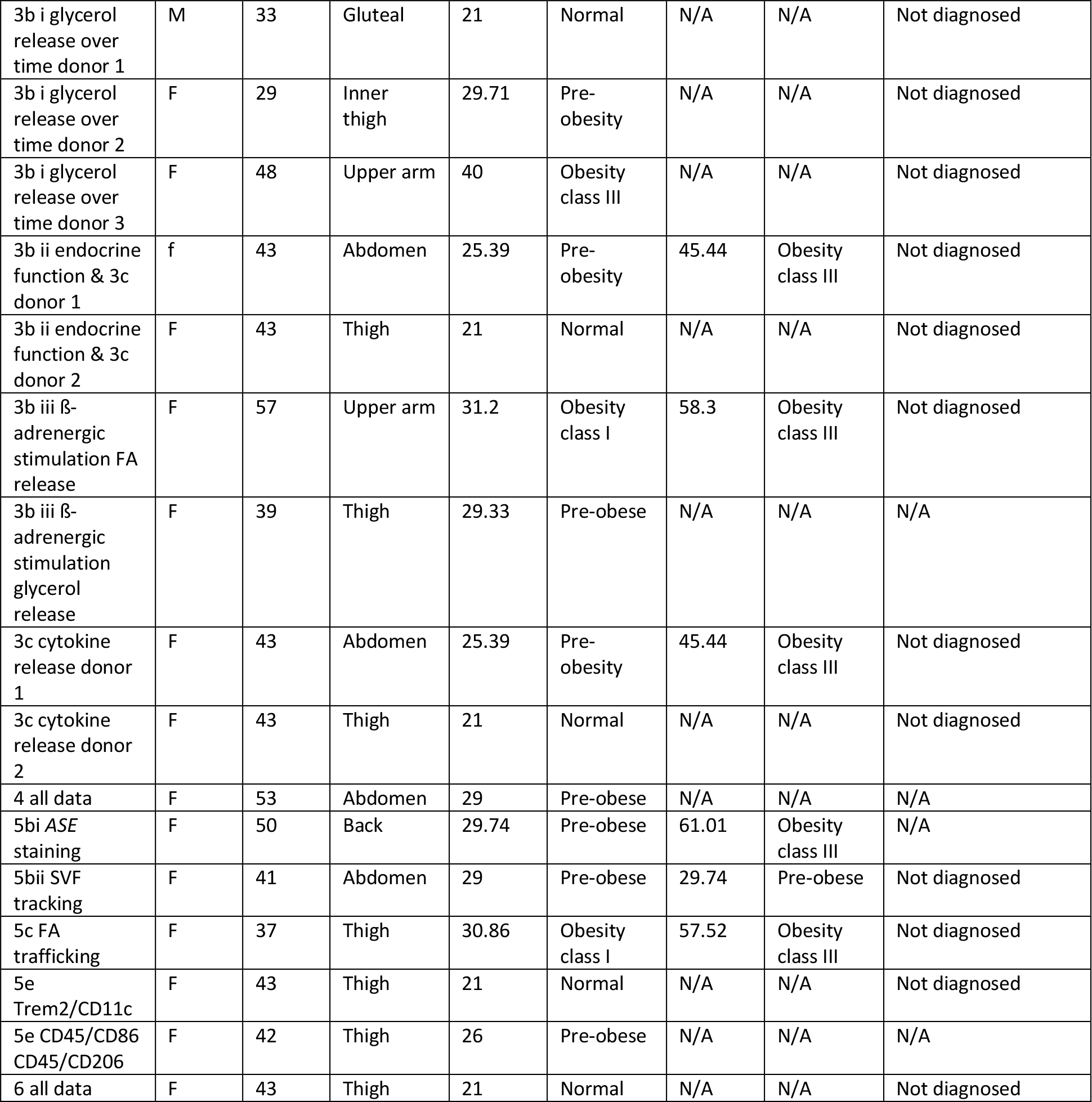
Overview of cell source and donor demographics itemized per experiment.

| Figure | Donor sex | Donor age | Source of biopsy | BMI at time of surgery | Weight status according to WHO | BMI at highest | Weight status according to WHO | Diagnosed metabolic disease (incl. diabetes) |
| --- | --- | --- | --- | --- | --- | --- | --- | --- |
| 3a morphology | F | 39 | Thigh | 29.33 | Pre-obese | N/A | N/A | N/A |
| 3b i MCFA/LCFA uptake | F | 30 | Abdomen | 23 | Normal | 44.9 | Obesity class III | Not diagnosed |
| 3b i glucose-dependent FA uptake | F | 57 | Upper arm | 31.2 | Obesity class I | 58.3 | Obesity class III | Not diagnosed |
| 3b i glycerol release over time donor 1 | M | 33 | Gluteal | 21 | Normal | N/A | N/A | Not diagnosed |
| 3b i glycerol release over time donor 2 | F | 29 | Inner thigh | 29.71 | Pre-obesity | N/A | N/A | Not diagnosed |
| 3b i glycerol release over time donor 3 | F | 48 | Upper arm | 40 | Obesity class III | N/A | N/A | Not diagnosed |
| 3b ii endocrine function & 3c donor 1 | f | 43 | Abdomen | 25.39 | Pre-obesity | 45.44 | Obesity class III | Not diagnosed |
| 3b ii endocrine function & 3c donor 2 | F | 43 | Thigh | 21 | Normal | N/A | N/A | Not diagnosed |
| 3b iii β-adrenergic stimulation FA release | F | 57 | Upper arm | 31.2 | Obesity class I | 58.3 | Obesity class III | Not diagnosed |
| 3b iii β-adrenergic stimulation glycerol release | F | 39 | Thigh | 29.33 | Pre-obese | N/A | N/A | N/A |
| 3c cytokine release donor 1 | F | 43 | Abdomen | 25.39 | Pre-obesity | 45.44 | Obesity class III | Not diagnosed |
| 3c cytokine release donor 2 | F | 43 | Thigh | 21 | Normal | N/A | N/A | Not diagnosed |
| 4 all data | F | 53 | Abdomen | 29 | Pre-obese | N/A | N/A | N/A |
| 5bi ASE staining | F | 50 | Back | 29.74 | Pre-obese | 61.01 | Obesity class III | N/A |
| 5bii SVF tracking | F | 41 | Abdomen | 29 | Pre-obese | 29.74 | Pre-obese | Not diagnosed |
| 5c FA trafficking | F | 37 | Thigh | 30.86 | Obesity class I | 57.52 | Obesity class III | Not diagnosed |
| 5e Trem2/CD11c | F | 43 | Thigh | 21 | Normal | N/A | N/A | Not diagnosed |
| 5e CD45/CD86 CD45/CD206 | F | 42 | Thigh | 26 | Pre-obese | N/A | N/A | N/A |
| 6 all data | F | 43 | Thigh | 21 | Normal | N/A | N/A | Not diagnosed |

The viability of adipocytes on-chip was assessed non-invasively via monitoring the release of lactate dehydrogenase (LDH) into the media effluents (supplementary figure S1). Therefore, effluents were collected every 24 h over a 12-day culture. While at the beginning of the culture low levels of LDH were detected (below 10% relative to expected maximum release), LDH was not detectable after d5 anymore. These findings indicated a good overall on-chip viability of adipocytes. Culture monitoring via bright field microscopy further backed the evidence of a stable adipocyte long-term culture on-chip (supplementary figure S2).

To characterize the morphology of adipocytes further, we stained lipid droplets (with a BODIPY neutral lipid stain), perilipin A (via immunofluorescence staining) and nuclei on d5 of on-chip culture (figure 3a). Confocal imaging of this staining revealed a dense, 3-dimensional arrangement of adipocytes throughout the entire chamber. Moreover, it confirmed the preservation of key morphological features of adipocyte maturity such as (i) unilocularity (i.e., storage of lipid content in one larger lipid vacuole instead of several smaller lipid vacuoles) and (ii) expression of the lipid droplet-coating protein perilipin A.^1^

**Figure 3.**
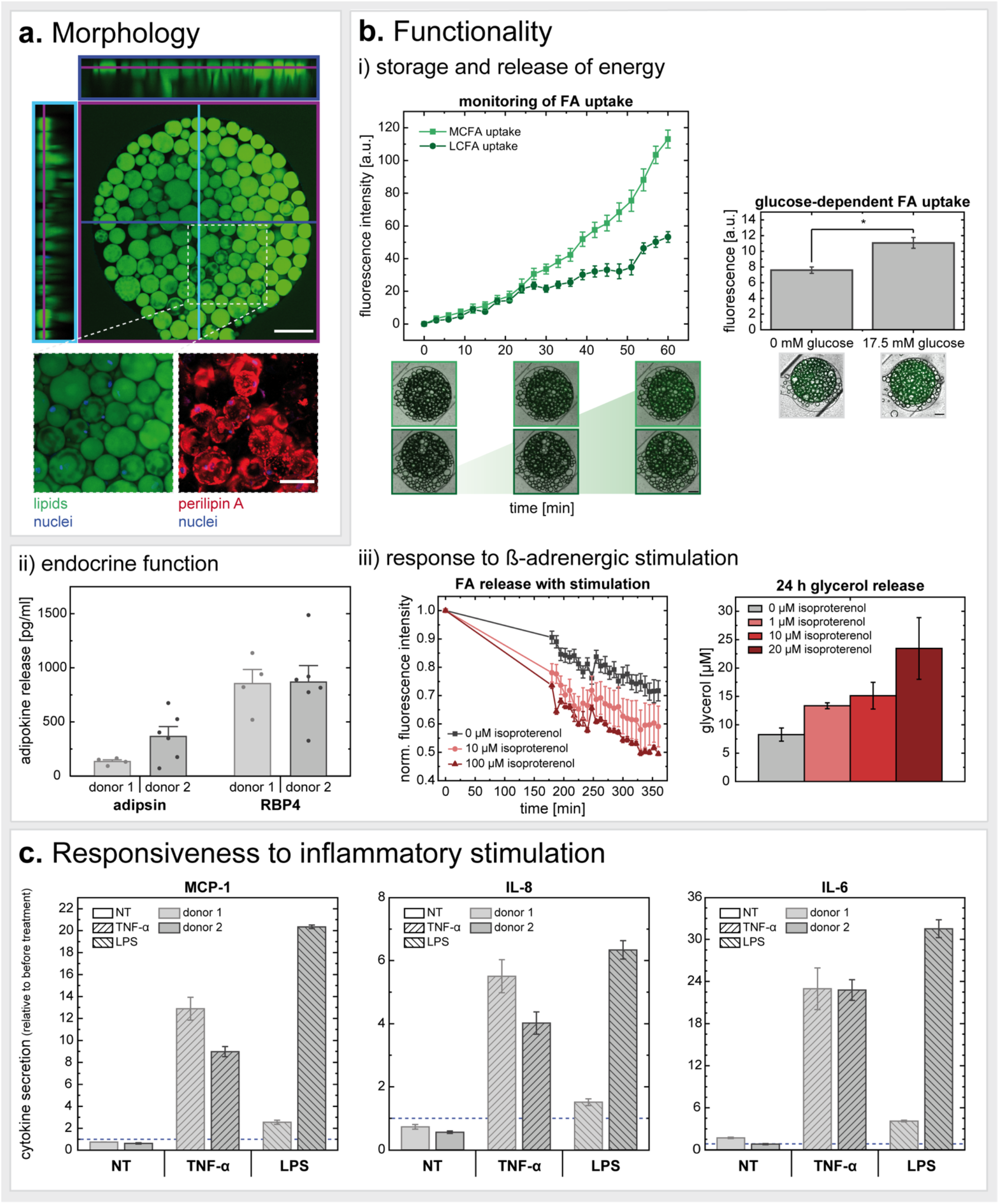
Characterization of on-chip human mature adipocytes. (a) On-chip visualization of mature adipocytes (fixed on d5) confirmed (i) 3-dimensional distribution of adipocytes inside the chips’ tissue chambers, and preservation of (ii) lipid content unilocularity as well as (iii) adipose-specific markers such as perilipin A. Scale bars equal 200 µm (orthogonal view) and 100 µm (maximum intensity projection of zoom-in/visualization of perilipin A). (b) For functional validation, we assessed (i) basal energy storage and release properties by monitoring uptake of medium- and long-chain fatty acid (FA) analogs (on d12) (MCFA n=14; LCFA n=16) and its dependency on glucose (on d4-d5) (no glucose n=3; high glucose n=6) (i). We further analyzed (ii) basal adipokine secretion (on d4) (donor 1 n=4; donor 2 n=6) as well as (iii) the adipocytes’ response to ß-adrenergic stimulation (on d4-d5). (c) Cytokine release in response to proinflammatory stimulation for 24 h with TNF-α (20 ng/ml) or LPS (100 ng/ml) on d5. Cytokine secretion is depicted relative to the secretion determined for the 24 h-period before stimulation (two independent chips per donor for each condition).

Unilocularity is a vital hallmark of the mature adipocyte phenotype. Many mature adipocyte *in vitro* culture methods, such as different variants of ceiling cultures (Sugihara et al., 1989; Zhang et al., 2000; Fernyhough et al., 2004), eventually induce a dedifferentiation of adipocytes to fibroblast-like progenitor states. Along the dedifferentiation process, the adipocytes undergo intracellular reorganization such as loss of the large lipid droplet, instead being multilocular, and spreading of cytoplasm (Côté et al., 2019). The dedifferentiation might be induced by an exposure to physical stressors (Yiwei Li et al., 2020), such as the presence of an adhesion surface as in the case of ceiling culture. The readiness of mature adipocytes to dedifferentiate has high potential for regenerative medicine (Côté et al., 2019), and elucidation of the underlying mechanisms of de- and redifferentiation is of utmost importance for understanding tumor progression (Yiwei Li et al., 2020, p.). Yet, this change in cell identity would be more than unfavorable when studying mechanisms of adipose tissues. Perilipin A, also called PLIN1, is expressed abundantly in mature adipocytes. It functions as a stabilizer of larger lipid droplets (usually < 10 µm) and plays an important role in hormone-induced lipolysis (Itabe et al., 2017). More recent studies in mice even suggest a major contribution of PLIN1 to anti-inflammatory processes and prevention of insulin resistance by restricting uncontrolled lipolysis (Sohn et al., 2018).

Adipocyte function on-chip was confirmed by analyzing its energy storage and mobilization capacities (figure 3bi). Upon administering fluorescently tagged fatty acids (FAs) to the adipocytes via the media perfusion, FAs were taken up by the adipocytes as indicated by an increase in intracellular fluorescence intensity. This uptake was monitored in real-time by imaging the individual tissue chambers every 3 minutes and quantified by plotting the fluorescence intensities against time of FA administration. Both fluorescent analogs of dodecanoic acid, also called lauric acid [with 12 carbon (C) atoms a representative of a medium-chain fatty acid (MCFA), BODIPY-C_12_], and hexadecanoic acid, also called palmitic acid [with 16 C atoms a representative of a long-chain fatty acid (LCFA), BODIPY-C_16_], were administered to capture potentially different FA uptake mechanisms. While short-chain fatty acids and MCFAs can freely diffuse across the cell membrane into the cytosol, the uptake of LCFAs, which are the most abundant among the three FA types, appears to be more complex (Schönfeld and Wojtczak, 2016). Despite still being under discussion, LCFA uptake might be realized through combination of passive diffusion and protein-accelerated entry into the membrane as well as desorption at the inner side of the membrane (Glatz and Luiken, 2020; Jay et al., 2020; Thompson et al., 2010).

Of note, the MCFA analog has two fluorophores attached (in positions C1 and C12), as compared to the LCFA analog, which has the BODIPY-fluorophore only in the C16 position. Therefore, the higher final fluorescence intensity signal obtained when feeding the MCFA analog could be attributed to the double amount of fluorophore. It is noteworthy that through the attachment of these two fluorophores, the MCFA analog might be comparable to a LCFA regarding its size. Hence, its trafficking properties could resemble that of a LCFA as well (Kolahi et al., 2016).

Leveraging the established FA uptake assay, we further investigated the dependency of BODIPY-C_12_ uptake rates on glucose concentration in the perfused medium. We found FA uptake rates to be higher, when the medium contained a high glucose concentration (17.5 mM) as compared to medium with no glucose added (except for glucose contained in fetal calf serum). These findings are in line with the need for glucose to form glycerol-3-phosphate for the backbone of triacylglycerides (TAGs) in lipogenesis (Morigny et al., 2021). Another reason might be the inhibition of fatty acid release by glucose uptake (Wolfe, 1998).

Furthermore, basal lipolytic activity during on-chip culture was determined by measuring glycerol concentration in media effluents for three different adipocyte donors. While a release of glycerol was detected for all donors, we found considerable inter-donor variations concerning the released concentrations (supplemental figure S3). These variations were found to be higher than intra-donor variations on different days of analysis as well as variations from different independent chips of the same donor on the same day of analysis.

This donor-specific cell behavior was also present when we determined basal adipokine release from the on-chip adipocytes (figure 3bii). While the release of retinol binding protein 4 (RBP4) was similar between the two donors, the secretion of adipsin varied. Generally, the adipokine release by the adipocytes on-chip demonstrates an endocrine functionality in addition to the metabolic functionality. Interestingly, adiponectin and leptin release into media effluents was not detectable for our adipocyte-only on-chip cultures; in co-culture with other WAT cell components, however, the release of these two important adipokines could be verified (cf. Autologous full complexity WAT-on-chip). Of note, adipose endocrine signaling occurs not only through peptides, such as adipokines and other cytokines but also through fatty acids (‘lipokines’) as well as exosomal microRNAs (Morigny et al., 2021; Scheja and Heeren, 2019).

The findings from our adipocyte-only chip culture experiments indicate that the platform is well suited for the culture of this demanding cell type. Adipocyte buoyancy and fragility is managed by encapsulation in a hydrogel matrix for cell anchorage and by very gentle injection and culture properties (sequential injection and protection from shear). Through a range of assays, we could show that a mature adipocyte phenotype as well as key *in vivo* functions were preserved in our *in vitro* model. Importantly, our on-chip culture concept was able to capture inter-donor differences concerning general adipocyte function, which could also be observed when studying responsiveness to external stimulations.

Next, we sought to study the adipocytes’ drug responsiveness. Due to its lipolytic effects, we selected the ß-adrenoreceptor agonist isoproterenol and administered 1-100 µM via the media perfusion (figure 3biii). As other catecholamines, this synthetic noradrenaline-derivative induces the breakdown of TAGs and its release from adipocytes. When introducing the drug after feeding the tissues with the BODIPY-C_12_ FA, we observed different FA release rates. The higher the isoproterenol concentration, the faster the intracellular fluorescence intensity signal from the BODIPY-C_12_ decreased. Another readout backing the adipocytes’ lipolytic response to ß-adrenergic stimulation was the determination of glycerol secretion during a 24 h drug treatment, which revealed a dose-dependent response; higher glycerol levels associated with higher doses of the drug.

Finally, we evaluated the adipocytes’ proinflammatory response to an acute 24 h TNF-α or LPS stimulation (figure 3c). We observed an increase in monocyte chemoattractant protein-1 [MCP-1, alternatively CC-chemokine ligand 2 (CCL2)], interleukin-8 [IL-8, alternatively C-X-C motif chemokine ligand 8 (CXCL8)] and interleukin-6 (IL-6) secretion for both TNF-α and LPS treatment. These findings were expected since adipocytes are responsive to both TNF-α and LPS, and have been shown to produce any of the three analyzed cytokines (Bruun et al., 2001; Hoch et al., 2008; Meijer et al., 2011). Again, we performed this experiment for two different adipocyte donors and found considerable differences in the degree of response as well as between the two stimulants. We further investigated the impact of inflammatory stimulation on adipocytes’ FA uptake as well as glycerol release; using the abovementioned methods, no difference in the examined properties were registered (data not shown).

### Characterization of on-chip endothelial barrier from mvECs

Next, we established protocols to line the perfused media channels, particularly the membrane forming the interface to the tissue chambers, with endothelium [microvascular endothelial cells (mvECs)]. Through this step, the transport from the media channel across the membrane into the tissue compartment was upgraded from passive diffusion to dynamic transport regulated by endothelial cells.

To characterize this on-chip endothelial barrier, we ran a series of experiments with endothelial barrier-only chips, i.e., no other cell type included in the tissue chambers. However, to maintain on-chip mechanical properties, the tissue compartment was filled with the hydrogel matrix prior to mvEC seeding into the media channels. After seeding, the cells were allowed to adhere overnight (under static, diffusion-driven nutrient supply) before connecting the chips to constant media perfusion. The flow rate was then ramped up to the final flow rate (20 µL/h) in a stepwise manner (increase by 5 µL/h every 2 h) to avoid mvEC detachment.

The mvECs quickly formed a dense monolayer that remained viable for at least one week of on-chip culture (figure S4a). We confirmed endothelial identity by verifying the expression of CD31 [alternatively platelet endothelial cell adhesion molecule 1 (PECAM1); a junctional molecule highly expressed on the surface of ECs] (figure 4a) as well as CD309 [alternatively vascular endothelial growth factor receptor 2 (VEGFR-2) or kinase insert domain–containing receptor (KDR)] and endothelial nitric oxide synthase [eNOS; alternatively nitric oxid synthase 3 (NOS3)] (figure S4b). Besides confirming EC identity, the anti-CD31 staining also demonstrates the formation of tight endothelial barriers with pronounced intercellular junctions throughout the entire chip. CD31 functions as a mechano-sensor, controls leukocyte trafficking and maintains the integrity of EC junctions (Privratsky and Newman, 2014). However, we did not observe any alignment of the mvECs in the direction of flow. This can be attributed to the shear forces on the membrane being considerably lower than *in vivo* (figure 2c) [here ∼ 3×10^-3^-4×10^-3^ dyn/cm^2^, figure 2c, *in vivo* usually 0.1–60 dyn∕cm^2^ (Park et al., 2011)]. An increase in flow rate was not possible: a flow rate higher than 20 µL/h diluted metabolites and messenger molecules secreted by the adipocytes into the effluent media to a concentration below detection limits of determination assays used in this study.

**Figure 4.**
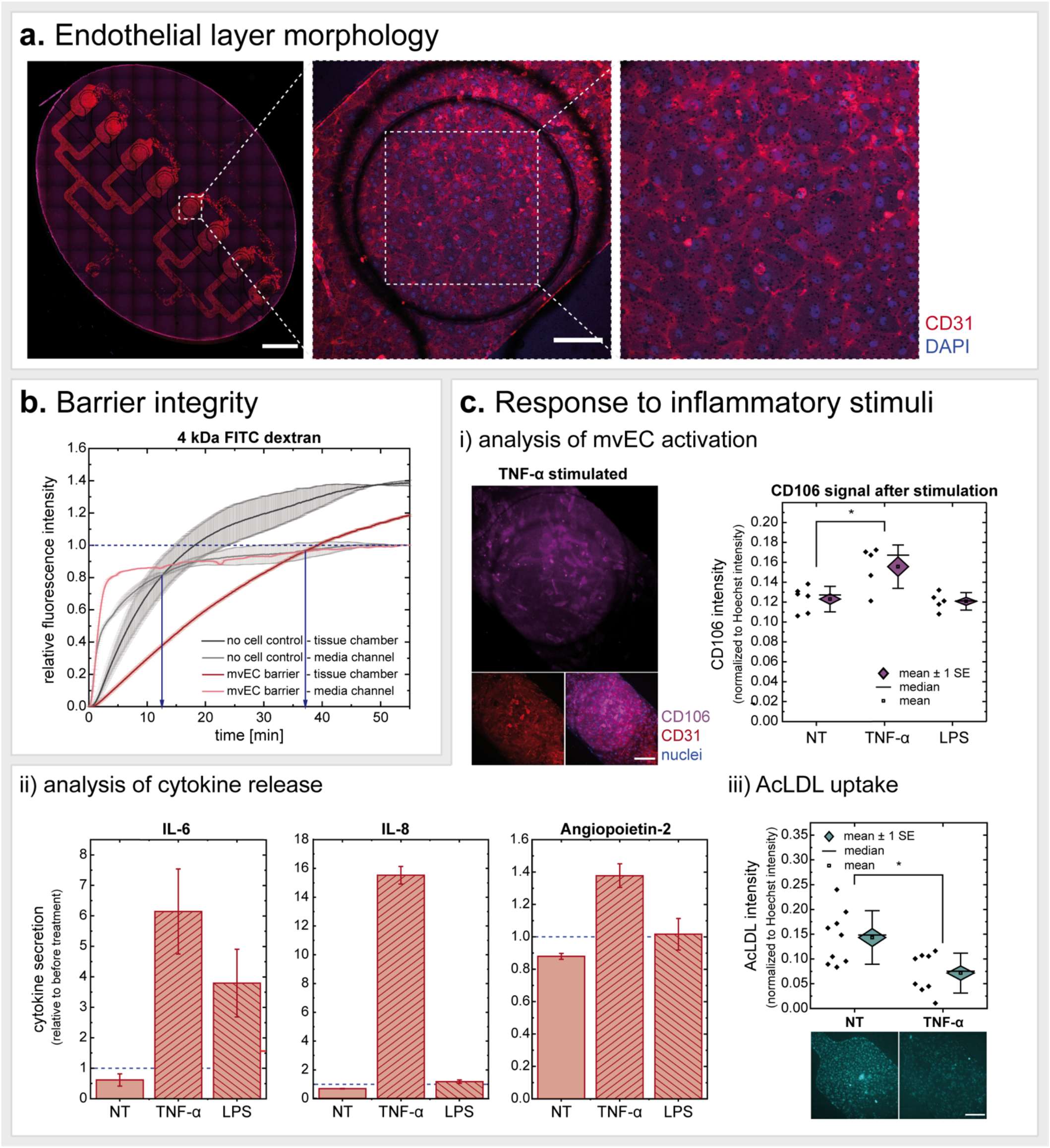
Characterization of on-chip endothelial barrier. (a) Microvascular endothelial cells (mvECs) seeded onto the membrane in the medium channel formed uniform, tight monolayers as visualized by CD31 staining (fixed on d7). Scale bars equal 2 mm (tile scan of entire chip) and 200 µm (one-chamber view). (b) Endothelial barrier integrity determined by fluorescence macromolecule tracing. We measured time difference in 4 kDa FITC-dextran signal equilibria in tissue chambers vs. media channels for chips with endothelial barrier vs. chips without endothelial barrier. Endothelial barriers were less permeable than plain, acellularized membranes. (c) Exposure of the on-chip endothelial barrier to inflammatory stimuli lead to (i) a significant increase in CD106 expression for TNF-*α* stimulation (unpaired *t* test with *p*-value ≤ 0.05; scale bar equals 200 µm), (ii) altered inflammatory cytokine secretion (n=2 for each stimulus) and (iii) decreased uptake/intracellular retention of acetylated low-density lipoprotein (AcLDL) (unpaired *t* test with *p*-value ≤ 0.05; scale bar equals 200 µm).

The permeability of the mvEC barrier was tested by analyzing the transport of a fluorescently labeled dextran across the endothelial barrier in comparison to an acellular membrane (figure 4b and figure S4c). We found that 4 kDa FITC-dextrans (figure 4b) are retained longer in the medium channel in the presence of a vascular barrier; in chips without this barrier, an equilibrium in fluorescence signal between media channel and tissue chamber was reached notably faster. Repeating this permeability assay using 40 kDa and 150 kDa FITC-dextrans suggested similar differences in macromolecule transport across the endothelial barrier (figure S4c).

Finally, we investigated the response of the endothelium to pro-inflammatory stimuli (figure 4c). We found, that the CD106 [alternatively vascular cell adhesion molecule 1 (VCAM-1)] expression by mvECs was significantly increased in response to TNF-*α*, but not to LPS. Compliant with this result, we found the endothelial release of proinflammatory cytokines (IL-6, IL-8 and angiopoietin 2) after stimulation to be most increased after the TNF-*α* treatment. While the LPS treatment induced an increased release of IL-6, releases of IL-8 and angiopoietin 2 were minimally increased by the proinflammatory challenge. Moreover, TNF-*α* stimulation affected the uptake (or intracellular retention) of acetylated low-density lipoprotein (Ac-LDL) by the endothelial cells resulting in almost twice as high fluorescence intensity.

Overall, these findings show that a tight endothelial barrier could be successfully established on-chip and key endothelial functions maintained for at least one week of culture. The vasculature is a dynamic barrier that can rapidly respond to changes in the circulation. Amongst other ways of such response, ECs take the part of metabolic gate keepers. They regulate and adjust transport rates of nutrients and hormones, including FAs, glucose and insulin, from the vessel lumen into tissues (Hasan and Fischer, 2021). Another important aspect of the endothelial barrier is its role as traveling route for immune cells and, potentially, recruiters thereof.

Besides the systemic endothelial tasks, ECs also contribute to organ-specific functions depending on their site of operation. Generally, owing to the adipose tissue’s enormous plasticity, adipogenesis is strongly dependent on angiogenesis (Augustin and Koh, 2017). Yet, the quiescent endothelium is as important as the active one: Adipose mvECs, for instance, were shown to directly crosstalk with adipocytes to regulate peroxisome proliferator activated receptor γ (PPARγ) pathways and thereby the adipocytes’ ability to take up and store lipids (Gogg et al., 2019). Notably, even though adipose mvECs take up lipids as well, they cannot undergo a full adipogenic differentiation when exposed to cues of adipose differentiation (Gogg et al., 2019). In this project dermal instead of adipose-derived mvECs were used owing to the limited amount of subcutaneous adipose tissue, which had to serve as a source for adipocytes, general SVF as well as tissue-resident immune cells. These dermal mvECs were isolated from the dermis of the skin/fat biopsy. Yet, despite the difference in origin tissue, it was shown previously that skin-derived and adipose-derived mvECs are very similar: they showed the same expression of endothelial markers, migration and sprouting behavior as well as response to inflammatory stimulation (Monsuur et al., 2016).

### Autologous full complexity WAT-on-chip: fit-for-purpose mix-and-match toolbox

To further enhance the physiological relevance of our model, we sought to integrate stromovascular cells in addition to the adipocytes and endothelial barrier. The SVF is a dynamic, heterogeneous cell population with variable degrees of maturity and varying functions. It is the sum of all adipose tissue nucleated cells except for adipocytes themselves, and it includes mesenchymal stem cells, adipocyte and vascular progenitors, mature vascular cells as well as fibroblasts and various types of immune cells. Since the cellular composition was reported to change greatly when culturing and passaging SVF (Nunes et al., 2013; Sun et al., 2019; Szöke et al., 2012), we injected the cells together with the adipocytes the day after isolation.

Notably, combining patient-specific adipocytes and mvECs presented a bigger logistical challenge: while adipocytes cannot be cultured properly in conventional cell culture formats and need to be injected into the microfluidic platform within 24 h after isolation, the mvEC yield after isolation is not sufficient for prompt injection. Instead, mvECs need to be expanded and purified from other cell types, such as fibroblasts, in adherent cell culture for at least seven days. Therefore, adipocytes were (co-)cultured with or without SVF/CD14^+^-cells on-chip for one week before ECs were set for seeding on d7 (figure 7, Materials and Methods section).

We created a physiologically relevant, autologous *in vitro* model of human WAT by integrating adipocytes, SVF and mvECs, all derived from the same tissue donor. To provide a fit-for-purpose model that allows researchers to choose the cell types of interest and the level of complexity depending on the actual research question, we established the following co- and multi-culture models (figure 5a): adipocyte-only systems (culture condition hereinafter dubbed “*A*”), adipocyte-endothelial co-culture systems (“*AE*”), adipocyte-SVF multi-culture systems (“*AS*”) and adipocyte-SVF-endothelial multi-culture systems (“*ASE*”). Due to the high importance of adipose-immune interactions, we further established a method to build adipocyte-CD14^+^ cell co-culture systems (“*AM*”) to enable directed studies on adipocyte-monocyte/macrophage crosstalk. To characterize the various co- and multi-culture WAT models, we analyzed on-chip viability, physiological structure, cytokine release as well as FA trafficking properties.

**Figure 5.**
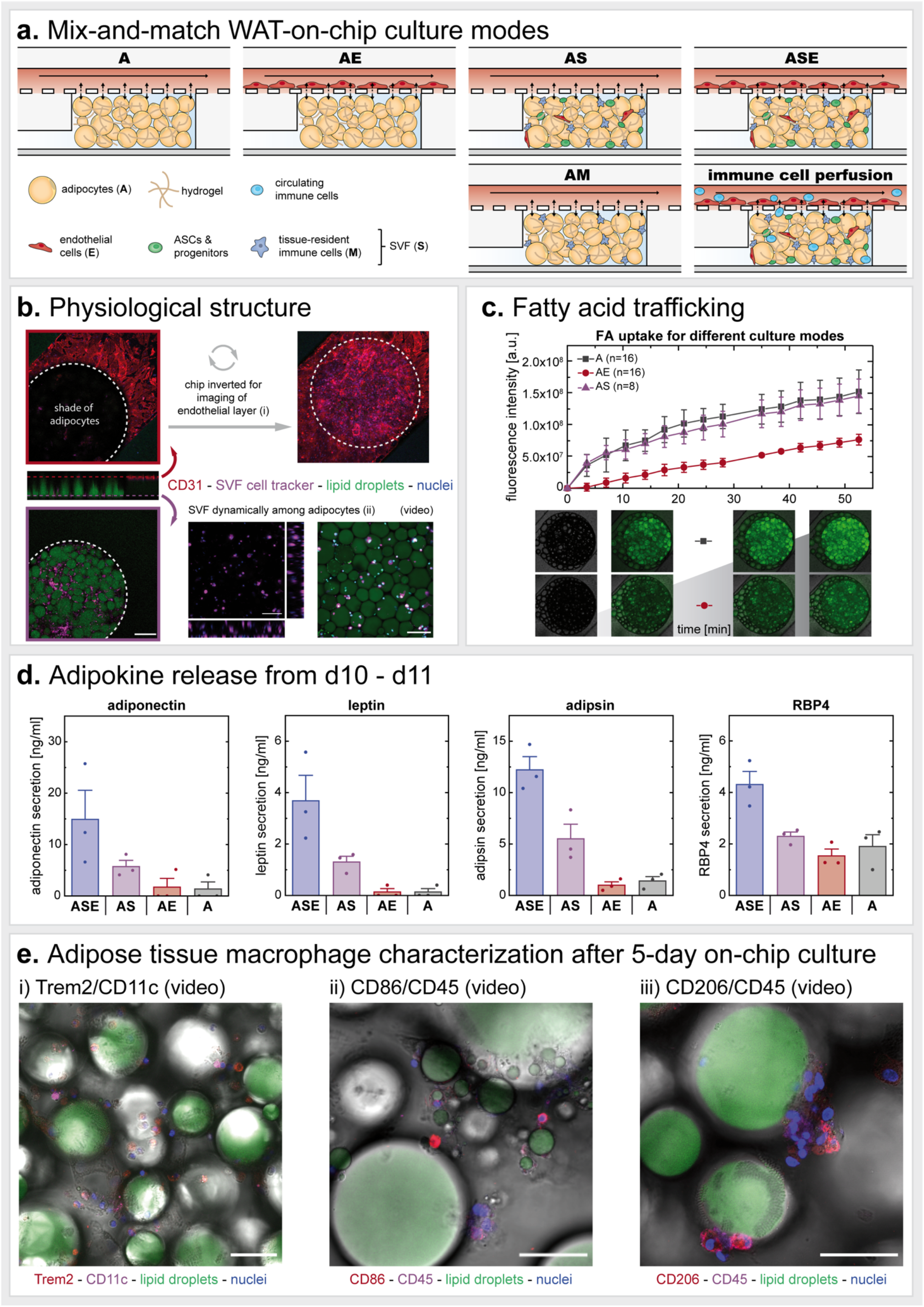
Modular mix- and-match toolbox to build fit-for-purpose autologous WAT-on-chip models. (a) To create WAT-on-chip models specifically tailored to certain scientific questions, we propose a spectrum of on-chip culture conditions with varying degrees of complexity ranging from simple adipocyte-only systems (*A*) to highly complex full WAT-on-chip models (*ASE*). (b) Structural characterization of the full WAT-on-chip model (*ASE*) on d12. Anti-CD31 immunofluorescence staining showed a tight endothelial barrier on the chip’s membrane. To visualize the membrane area over the adipocyte-filled tissue chambers, the chip had to be inverted (i). Adipocytes were displayed by staining of their lipid droplets. To uncover stromovascular cells, SVF was labelled with a cell tracker prior to injection into the chip. SVF was 3-dimensionally distributed among the adipocytes in the tissue chamber (adipocytes not shown for better visibility of SVF) (ii; orthogonal view). Moreover, a tracking of SVF motion within the first 6 h after injection revealed dynamic migration for some of the labelled cells while others remained stationary (ii; video). Scale bars equal 200 µm (one-chamber view) and 100 µm (zoomed-in orthogonal view and video). (c) Monitoring of FA trafficking properties for *A*, *AS* and *AE* systems uncovered noticeable differences in FA uptake comparing *A* and *AS* to *AE*. Representative images of *A* and *AE* conditions at time points 0 min, 10.5 min, 38.5 min and 52.5 min. (d) Comparison of adipokine release by different co-/multi-culture WAT-on-chips from the same patient, measured from media effluents collected for 24 h from d10 to d11 of on-chip culture. Even though the analyzed adipokines are exclusively produced by adipocytes, there were considerable differences in their release regarding the models’ cellular composition. (e) Identification of on-chip ATMs by visualizing CD11c and Trem2 (i), CD86 and CD45 (ii) and CD206 and CD45 (iii) expression. Lipid scavenging properties and formation of crown-like structures were visualized. Z-stack imaging data are represented as videos to ensure full elucidation of all events in one stack. Frames of videos are individual planes of acquired Z-stacks. Scale bars equal 50 µm.

The platform was able to maintain good overall viability of all cell types for at least 12 days of on-chip culture, as demonstrated by low LDH release into the media effluents (figure S1).

To confirm a physiological tissue structure, we fluorescently stained and imaged an *ASE*-chip after 12 days of on-chip culture (figure 5b). Endothelial cells – marked by an anti-CD31 staining – formed an interconnected monolayer (figure 5bi), comparable to our endothelial-only experiments (figure 4a). Adipocytes in the *ASE* chip also displayed large unilocular lipid droplets indicating preservation of adipocyte maturity. Because of their heterogeneity, there is no single specific marker to identify SVF. The location of the stromovascular cells could, nevertheless, be monitored by labelling them with a cell tracker prior to injection, demonstrating a homogeneous distribution across the tissue chamber among the adipocytes (figure 5bii). Moreover, tracing of SVF migration on-chip for 6 h after injection revealed that some cells moved dynamically throughout the tissue chamber while most cells appeared to have settled in defined locations, pointing to the heterogeneity in function (figure 5bii video).

Characterization of FA transport properties via the BODIPY-C_12_ FA uptake assay showed similar uptake rates between *A* and *AS* conditions. The *AE* systems, however, displayed a noticeably (approx. 50%) lower fluorescence intensity (figure 5c), hinting at a lower FA uptake and/or actively controlled transport through the EC barrier. From the results of the EC barrier integrity characterization using a 4 kDa FITC-dextran (10x bigger than the BODIPY-C_12_ FA analog; figure 4b), a convergence of fluorescence intensity signals would have been expected within the duration of the assay, assuming comparable intracellular FA uptake by adipocytes across the three conditions and merely passive transendothelial transport of the BODIPY-C_12_ FA analog. Yet, after 50 min, the fluorescence intensity signal for *AE* was still at approx. 50% of the one for *A* and *AS*. This suggests that either the FA uptake is downregulated by the interaction of ECs and adipocytes or that the endothelial-mediated FA transport is not passive, diffusive transport but an actively controlled process. Indeed, in its function as metabolic gatekeeper, the endothelium is presumed to actively adjust its barrier function in order to regulate FA and lipoprotein transport by involving a complex signaling machinery (Mehrotra et al., 2014; Pi et al., 2018). Furthermore, in line with previous reports (Gogg et al., 2019), the increased background fluorescence signal for the *AE* condition, as compared to *A*/*AS,* indicates an uptake of FAs by ECs.

Next, we compared the adipokine (adiponectin, leptin, adipsin and RBP4) release from four different culture systems (figure 5c). Interestingly, the *ASE* condition showed the highest release for all four adipokines. On the contrary, for *AE* and *A,* adipokine release was overall much lower than for *ASE*: adiponectin and leptin release was hardly detectable while adipsin and RBP4 levels were only approx. 8-10% and 35-45%, respectively. This outcome highlights how crucial the contribution of the other cell populations in adipose tissue is to adipocyte function and how important it is to integrate them to model adipose tissue: each of the analyzed factors has been reported to be predominantly produced by adipocytes (except for RBP4, which is also produced by hepatocytes); no other adipose-associated cell type was described to release any of the four cytokines in significant amounts (Scheja and Heeren, 2019). Hence, given the four adipokines are indeed exclusively secreted by the adipocytes, the increase in secretion for the *ASE* condition stems from the interaction of the adipocytes with the other cell types.

Finally, we established an adipose-on-chip model that integrates solely adipocyte and macrophages (*AM*) amenable for studies specifically targeted at the interaction of these two cell types. We visualized immune cell phenotypes on d5 after injecting a mixture of adipocytes and CD14^+^ cells isolated from patient-specific SVF (figure 5e). CD14 is a co-receptor to the LPS receptor and is strongly expressed on monocytes and macrophages. Adipose tissue macrophages (ATMs) exhibit a great phenotypic plasticity, that is much more complex than the binary M1/pro-inflammatory-vs. M2/anti-inflammatory classification and include populations such as metabolically activated ATMs (MMe) or oxidized ATMs (Mox) (Yunjia Li et al., 2020; Russo and Lumeng, 2018). Additionally, a new subset of lipid-associated macrophages (LAMs), highly expressing the lipid receptor Trem2, was found in adipose tissue from obese humans. Trem2 was discovered to be essential for LAMs to exert their protective functions such as counteracting adipocyte hypertrophy and inflammation. (Jaitin et al., 2019) Moreover, many macrophages have a mixed activation state and harbor both M1 and M2 markers, such as the hybrid CD11c^+^ (classically M1) CD206^+^ (classically M2) ATMs associated with insulin resistance (Wentworth et al., 2010).

After five days of on-chip culture, we could observe ATMs in the tissue expressing the commonly occurring markers of human ATMs CD11c, CD206 as well as CD86, the general leukocyte marker CD45 as well as LAM-specific marker Trem2 (figure 5e). The ATMs were positioned in 3D among the adipocytes with clusters of ATMs frequently attaching to individual adipocytes or even wrapping around them as crown-like structures (e.g., video 5e). Visualization of lipid droplets via both neutral lipid staining and bright field microscopic imaging, moreover, revealed an uptake of lipids by ATMs indicating a lipid scavenging activity (e.g., video 5e). We also investigated monocyte/macrophage phenotypes under inflammatory stimulation: Via immunofluorescence staining, we did not observe any obvious differences in ATM marker expression between the stimulation conditions (LPS, TNF-*α*, NT). Effluent analysis with respect to secretion of pro-inflammatory cytokines, however, revealed an upregulation of several typical pro-inflammatory cytokines during TNF-*α* or LPS treatment (figure S5).

This capacity of the model to integrate ATMs is of particular importance, since with WAT’s recognition as an endocrine, immunoregulatory organ, adipose tissue immune cells, specifically ATMs, have become a prominent research topic. As described above, ATMs are extremely plastic and adapt to different adipose tissue physiological states. In the healthy state, they regulate tissue homeostasis while in diseased conditions, such as obesity, ATMs play a major role in the low-grade, chronic inflammation and dysregulated metabolism (Caslin et al., 2020). Their tasks are as multifaceted as their appearance, but one of the main ATM functions seems to be engulfing dead adipocytes. Being under severe metabolic stress predisposes hypertrophic adipocytes to pyroptosis (i.e., a pro-inflammatory form of programmed cell death), a process attracting macrophages (Giordano et al., 2013). Aside from ingesting entire cells, ATMs are attributed to fulfill substrate buffering activities: Given their ability to handle variable substrate loads throughout their lifetime, ATMs can incorporate lipids, catecholamines and iron to modulate the availability of and protect other cell types from a surplus of these substances in the adipose tissue microenvironment (Caslin et al., 2020; Caspar-Bauguil et al., 2015).

There is also evidence for an active, direct crosstalk between adipocytes and ATMs. Adipocyte-derived FAs are an important modulator of macrophage metabolism; for instance, ATM uptake of FAs was found to be coupled to lysosome biogenesis (Xu et al., 2013). Moreover, adipocytes were found to release exosome-sized, lipid-laden vesicles (AdExos) that were found to induce a differentiation of macrophage precursors into ATMs. Hence, AdExos might not only be an alternative lipid release mechanism from adipocytes but also a directed technique for adipocytes to modulate macrophage function. (Flaherty et al., 2019) Another important example of adipocyte-macrophage crosstalk is the recently discovered intercellular transfer of mitochondria: in an *in vivo* rodent study, mitochondria were found to be transferred from adipocytes to neighboring macrophages, potentially to support the survival of metabolically compromised cells (Brestoff et al., 2021).

While adipocytes’ dysregulated metabolism still appears to be the main drivers of WAT immune responses (Morigny et al., 2021), ATM contributions should not in the least be disregarded; their close and manifold crosstalk with the adipocytes still makes them key players of all adipose-associated pathologies and therefore a potential therapeutic target. Here, we have shown that our WAT-on-chip model can incorporate ATMs that maintain physiological phenotypes even after five days of on-chip culture and recapitulate key ATM functions such as scavenging lipids and phagocyting adipocytes and thereby serve as a powerful platform to further enlighten adipocyte-ATM interactions.

### Immunocompetency of the WAT-on-chip

After verification of the WAT-chip’s suitability to integrate macrophages, we expanded experiments on immunocompetency by elucidating immunomodulatory cytokine release and responsiveness to inflammatory threats. As already addressed, on-chip adipocytes and endothelial barriers respond to TNF-*α* as well as LPS stimulation by ramping up their release of pro-inflammatory cytokines (figures 3c and 4cii). Here, we examine the participation of (i) the other adipose tissue components and (ii) intercellular crosstalk in immune responses. Intriguingly, we already registered differences in basal (i.e., unstimulated) cytokine release between the different culture modes although all cells for all modes were derived from the same donor to maximize comparability (figure 6a). Generally, cytokine expression was lowest from adipocyte-only chips, indicating major contributions of the other adipose tissue-associated cells. While the MCP-1 concentration was comparable between *ASE*, *AS* and *AE*, IL-8 and IL-6 concentrations were highest for *AS*. It is surprising that the interleukin secretion from *ASE* is lower than the secretion from *AS*, since *ASE* contains the same number of adipocytes and SVF as the *AS* condition. This points to a damping impact of EC presence regarding IL-6 and IL-8 release into the medium compartment in the *ASE* condition and might be an example of the necessity for a holistic view when modeling intercellular communication. Overall, the measured cytokine concentrations fall in the same range as determined for other *in vitro* studies. WAT is a main source of circulating IL-6, with an about 35% contribution to basal circulating IL-6 (Wueest and Konrad, 2020). Besides adipocytes, adipose-derived mesenchymal stem cells were previously found to produce loads of IL-6, too (Blaber et al., 2012); hence, the elevated IL-6 release for the *ASE* and *AS* on-chip conditions is not surprising. While IL-6 has a context-dependent role and can act both pro- and anti-inflammatory, IL-8 and MCP-1 secretion is usually associated with inflammation and obesity (Cimini et al., 2017; Kim et al., 2006). Surprisingly, we still detected a basal release of both cytokines in the supposedly ‘healthy’ condition, which could lead back to the tissue origin – fat-removal procedures often occur in case of donor obesity. However, when adding external inflammatory stimuli their secretion was massively (up to 70-fold) increased (figure 6b). Hence, it remains obscure whether the registered ‘basal’ cytokine release is already, to some extent, shifted into an inflammatory state due to an obesogenic ground state of donor cells, or whether IL-8 and MCP-1 are released from adipose tissue in moderation in a healthy state as well.

**Figure 6.**
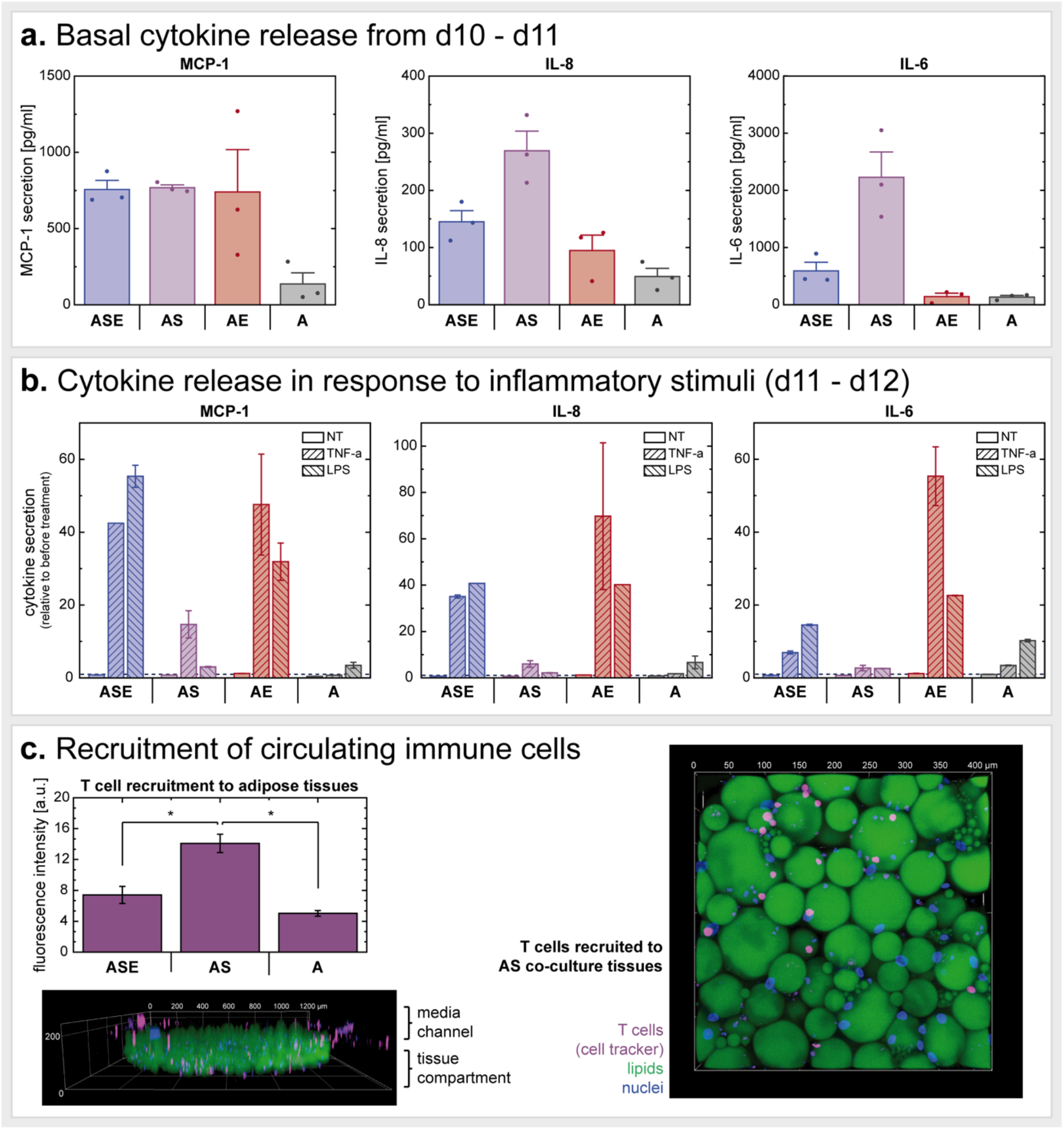
Immunocompetency of different WAT-on-chip culture conditions. (a) Baseline (i.e., non-stimulated) cytokine release from culture conditions *ASE*, *AS*, *AE* and *A* measured for 24 h from d10-d11. (b) Cytokine release in response to pro-inflammatory challenge (with TNF-*α* or LPS) for 24 h from d11-d12 on-chip. Cytokine concentrations are presented relative to the respective baseline cytokine release for each individual chip. (c) Recruitment of autologous T-cells perfused through the media channels for 18 h from d12-d13 of on-chip culture. Prior to perfusion, T cells were labeled with a cell tracker. Within the 18 h, T cells infiltrated the tissue chambers from the media channels, shown exemplarily in 3D renderings of an entire tissue chamber (side view) and of a zoom-in (top view). Recruitment was quantified by comparing fluorescence intensity in the tissue chambers (unpaired t test with p-value ≤ 0.05). Both fluorescence images show T-cell recruitment into *AS* chambers. All experiments were conducted simultaneously and with cells from the same donor.

As anticipated, in response to proinflammatory stimulation, cytokine release was upregulated in all culture conditions in comparison to non-treated systems (figure 6b). The increase of cytokine secretion relative to the 24 h period before treatment is highest for the *ASE* and particularly *AE* conditions implying a strong involvement of ECs to the inflammatory reaction. Moreover, the *ASE* and *A* chips appeared to respond stronger to the LPS treatment while *AS* and *AE* tended to release higher concentrations of the cytokines when stimulated with TNF-*α*. Notably, the temporal resolution of cytokine measurement in response to stimulation (here 24 h) could in the future be increased by sampling effluents, e.g., every 30 min to capture secretion kinetics.

Finally, we evaluated the suitability of our WAT-on-chip platform for studying immune cell perfusion and recruitment to adipose tissue. From d12-d13, autologous CD3/CD28 co-stimulated and fluorescently labelled T cells were perfused through the media channels of different adipose tissue culture mode chips for 18 h (figure 6c). By confocal imaging, we could confirm the infiltration of cell tracker-labelled cells into the adipose tissue compartment. This analysis indicated a significantly higher recruitment to the *AS* condition compared to both *ASE* multi-culture chips and *A* culture chips. A potential reason for this finding might be the elevated IL-8 and IL-6 release we registered from the *AS* cultures; both interleukins were shown to locally mediate T cell attraction (Bruun et al., 2001; McLoughlin et al., 2005; Taub et al., 1996; Weissenbach et al., 2004).

Additionally, we investigated the recruitment of PBMC-derived CD14^+^-cells to on-chip adipocytes upon perfusion through the media channels. While perfused CD14^+^-cells were not able to transmigrate through 3 µm pore-sized membranes, we were able to register scarce CD14^+^-cell infiltration into adipocyte-only systems when using 5 µm pore-sized membranes instead (figure S6). A potential reason for the low recruitment compared to T cell recruitment might be that the chemotactic cues produced by adipocytes were too low to attract monocyte-masses. However, these are just preliminary findings, and they would require more in-depth analysis.

Generally, these findings show that our platform is well suited to recapitulate T cell infiltration into adipose tissue. Importantly, it is not only the tissue-resident immune cells that seem to provide the necessary chemotactic cues; mature adipocytes themselves were able to attract T cells, too, which is in line with previous findings (Poloni et al., 2015). Incorporating T cells in adipose tissue models is of importance since T cells were recently discovered to play a major role in immunometabolism. WAT has been implicated in serving as a reservoir for tissue-resident memory T cells, which have distinct functional and metabolic profiles (Han et al., 2017). Furthermore, obesity has been associated with increased T cell populations, particularly adaptive CD8^+^ T cells, residing in WAT (Wang et al., 2021). Obesity might cause adipose tissue T-cell exhaustion (Porsche et al., 2021) and lead to unusual, T-cell mediated pathogeneses upon infection (Misumi et al., 2019).

## Conclusion

Here, we introduce, to the best of our knowledge, the first human, fully autologous immunocompetent WAT-on-chip platform integrating almost all in vivo WAT-associated cell types in mature states. Given the high complexity and associated logistical requirements, which accompany a full WAT in vitro model, we propose a mix-and-match WAT-on-chip toolbox that allows researchers to build a flexible, fit-for-purpose platform. As key component, mature human adipocytes are combined with patient-specific stromovascular cells, endothelium and/or different types of immune cells. The developed system enables long-term culture of human WAT in vitro while it preserves key functional features of not only adipocytes but at all other WAT-associated cellular components. More precisely, the system enabled a preservation of mature phenotypes alongside key responsibilities such as energy storage and mobilization functions, and basal endocrine activity. We further confirmed the on-chip WAT’s drug- and inflammatory responsiveness, and suitability for studies on immune cell infiltration.

The system is based on a specifically tailored microfluidic platform integrating several injection and on-chip culture features, such as the sequential loading and shielding from shear stress, that make it particularly favorable for the integration of human WAT. Thereby, the system is well equipped to overcome two major difficulties arising from working with human mature adipocytes: buoyancy and fragility. In the future, technological refinements could aim at scale-up and sensor integration: To achieve parallelized platforms with increased throughput, a combination with enabling technologies might be worthwhile; e.g., the WAT-on-chip design could be transferred to the organ-on-disc platform (Schneider et al., 2020; Stefan Schneider et al., 2021). To enable in-line monitoring of adipose tissue secretions, on-chip sensors based on different technologies could be integrated (Fuchs et al., 2021; Hu et al., 2020; Zhu et al., 2018, 2021).

To set up the WAT model, we established protocols and processes that allow to source all the different cell types from one individual donor; more specifically, from tissue samples of subcutaneous adipose tissue with skin that are readily available in most hospitals. The choice of a human cell source is an integral part for human-centered research. Even though findings from animal models have shed light into many aspects of WAT (patho-)physiology in the past, they are stretched to their limits regarding human-relevant mechanisms with increasing frequency. Despite efforts on humanization of animal models, there are still dominant discrepancies between rodents and humans, especially when it comes to studying metabolism and functioning of the immune system (Greek and Menache, 2013; Mestas and Hughes, 2004; van der Worp et al., 2010). Moreover, the capability to isolate all cell types from one individual, the bottom-up approach of tissue generation and the microscopic footprint of the WAT-on-Chip, enable not only fully autologous models but also the creation of a large number of tissue models from one donor; the latter allows circumventing the challenge of inter-donor differences as well as the study of patient-specific WAT responses, paving the way for future applications in personalized medicine.

When addressing patient-specificity, an aspect that is of particular interest is the immune system, which is closely interwoven with adipose tissue biology. Here, the autologous character of our WAT-on-Chip is of notable relevance especially when studying the adaptive immune system. The findings on WAT immunomodulatory functions on-chip back the suitability of our platform for research on adipose tissue inflammation and its underlying mechanisms. Especially the precisely controllable administration of inflammatory agents and the potential for highly time-resolved readouts of tissue response make our platform a powerful tool to study immune responses.

Besides demonstrating its competency to build a WAT-on-chip with highest physiological relevance integrating almost all cell types existing in in vivo WAT, we introduced a modular WAT-on-chip toolbox. When deciding on a suitable model for probing a specific scientific question, the rule should always be “as simple as possible but as complex as necessary”. In other words, the full WAT-on-chip model might be too complex for certain questions, and complexity might generate noise and off-target signals. The modular approach allows for an adjustable degree of complexity and enables end users to select those cellular components required for their specific question creating a fit-for-purpose, customized WAT-on-chip. For proof-of-principle, we characterized several combinations of mature adipocytes with stromovascular cells, ECs, and tissue-resident innate immune cells. Interestingly, we did not encounter any differences in adipocytes’ phenotype or energy storage function when comparing the different multi-culture conditions, but their endocrine function appeared to be strongly impacted by the co-culture with other cell types. Adipokines, presumably exclusively produced by adipocytes themselves, were found in highest concentrations in the full WAT model – notably higher than in the adipocyte-only condition. Furthermore, we found the response to proinflammatory threats modulated by influence of other adipose tissue-associated cells. Overall, the outcome of our study indicates that the full WAT model with all in vivo components indeed reflects adipocyte endocrine function best.

In conclusion, our novel WAT-on-chip system provides a human-based, autologous and immunocompetent in vitro model of WAT. It recapitulates almost full tissue heterogeneity by integrating not only mature adipocytes but also organotypic endothelial barriers and stromovascular cells, with optional separation of tissue-resident innate immune cells, specifically ATMs. Therefore, the new WAT-on-chip model can be a powerful tool for future, human-relevant research in the field of metabolism and its associated diseases as well as for compound testing and personalized- and precision medicine applications.

## Materials and methods

### Chip fabrication and characterization

#### Chip design and dimensions

The microfluidic platform used for integrating human WAT and associated immune components is a custom-made device consisting of two layers of micro-patterened polydimethylsiloxane (PDMS; Sylgard 184, Dow Corning, USA) sandwiching an isoporous, semipermeable polyethylene terephthalate (PET)-membrane (3 µm poresize: r_P_ = 3 μm; ρ_P_ = 8 × 10^5^ pores per cm^2^; TRAKETCH^®^ PET 3.0 p S210 × 300, or 5 µm poresize: r_P_ = 4.6 μm; ρ_P_ = 0.6 × 10^5^ pores per cm^2^; TRAKETCH^®^ PET 5.0 p S210 × 300, SABEU GmbH & Co. KG, Northeim, Germany). While the lower micro-patterned PDMS layer accommodates human WAT, the upper PDMS-layer, separated from the lower one by the membrane, serves as media compartment for constant media perfusion. To assure best optical accessibility of the tissues, the tissue compartment is secured to a glass coverslip (AN-21-000627; 25 mm x 75 mm, thickness 1, Langenbrinck GmbH, Emmendingen, Germany). The architecture of the microstructures in the PDMS layers was specifically designed to house human mature adipocytes, and it was drafted using CorelCAD [Corel Corporation, Ottawa, Ontario, Canada]. Table 1 and figure 2a provide an overview of the most important chip dimensions and resulting volumes.

#### Chip fabrication by soft lithography and replica molding

The microfluidic platforms were fabricated using the soft lithographic as well as replica molding protocols described in our previous publications (Loskill et al., 2017; Rogal et al., 2020, 2021). In brief, media channel- and tissue chamber microstructures in the PDMS layers were generated by using two differently patterned master wafers functioning as positive molding templates. These master wafers were produced by common soft lithography techniques first introduced by Xia and Whitesides (Xia and Whitesides, 1998). To achieve different heights for injection channels and tissue chambers in the tissue compartment layer, or a membrane inlay with media channel on top in the media compartment layer, respectively, each of the two master wafers was fabricated in two consecutive patterning steps. We recently described our procedures for a two-step master wafer fabrication in detail (Rogal et al., 2021). The PDMS layers were then generated by deploying two different replica molding techniques: (i) To create thin tissue layers with chambers and injection port structures opened to both sides, we used a technique called ‘exclusion molding’. By placing a release liner [Scotchpak™ 1022 Release Liner Fluoro-94 polymer Coated Polyester Film; 3M, Diegem, Belgia] onto the uncured PDMS on the wafer and applying uniform pressure onto the construct, curing of the PDMS resulted in 200 µm-high PDMS layers, as defined by the 200 µm tissue chamber height. Consequently, microstructures were open to both sides. (ii) The PDMS layer patterned with the media compartment was generated by standard replica molding; the amount of PDMS yielding a layer of approximately 5 mm was poured onto the master wafer and released after curing. Then, the PDMS slabs resulting from the molding processes were cut to the size of the chip and ports to access the chips were pierced using a biopsy punch (Disposable Biopsy Punch, 0.75 mm diameter; 504529; World Precision Instruments, Friedberg, Germany). To enable O_2_-plasma-based bonding of the PET-membrane, commercially available membranes were functionalized by a plasma-enhanced, chemical vapor deposition (PECVD) process (Rogal et al., 2020). In a three-step O_2_-plasma activation-(each 15 s, 50 W, 0.2 cm^3^m^-1^ O_2_; Diener Zepto, Diener electronic GmbH + Co. KG, Ebhausen, Germany) and bonding sequence, the chip was assembled: (i) bonding of exclusion-molded tissue compartment layer to a glass coverslip, (ii) bonding of functionalized PET-membrane into membrane inlay of media compartment layer, and (iii) full chip assembly by bonding the media compartment layer with membrane to the tissue compartment layer on the coverslip. To enhance bonding, the assembled chips were kept at 60°C overnight. To assure quality of bonding, chips were then flushed with DI-water and observed for any leakages or discontinuities in liquid flow. One day prior to cell injections, the chips were sterilized and hydrophilized by a 5-minute O_2_-plasma treatment. Under sterile conditions, they were then filled with PBS^-^ and kept overnight fully immersed in Dulbecco’s phosphate buffered saline without MgCl_2_ and CaCl_2_ (PBS-; Merck KGaA) to allow for any residual air to evacuate from the channel system.

#### Numerical modeling

To model fluid flow, its associated shear forces as well as transport of diluted species, we used COMSOL Multiphysics (COMSOL Vers.5.5, Stockholm, Sweden). The numerical model was based on simulations we previously published for our murine as well as our precursor adipocyte-on-chip models (Loskill et al., 2017; Rogal et al., 2020). In brief, we coupled the Free and Porous Media Flow and Transport of Diluted Species in Porous Media physics modules. We included the presence of hydrogel in the tissue compartment for all simulations since it significantly affects the convective and diffuse flow regimes. We used the Navier-Stokes equation with the properties of water (dynamic viscosity μ = 1 × 10^−3^ m^2^/s, density ρ = 1000 kg/m^3^) to model incompressible stationary free fluid flow at a general flow rate of 20 µl/h (equivalent to a flow rate of 2.5 µl/h in each of the eight parallel media channels over the tissue chambers). To model fluid flow from the media channel through the porous PET-membrane into the tissue chamber as well as through the hydrogel, Darcy’s law was used (membrane - porosity = 0.056, hydraulic permeability κ = 1.45 × 10^−14^ m^2^; hydrogel - porosity = 0.99, hydraulic permeability κ = 1.5 × 10^−16^ m^2^ (McCarty and Johnson, 2007)). Using the time-dependent convection-diffusion with diffusion coefficients of 1 × 10^−9^ m^2^/s (water) and 1 × 10^−11^ m^2^/s (hydrogel) and an initial concentration of 1 mol/m^3^, we described transport of diluted species.

### Isolation and culture of primary adipose tissue- and blood-derived cells

#### Human tissue samples

All cell types [i.e., adipocytes (‘*A*’), cells from SVF (‘*S*’), microvascular endothelial cells (‘*E*’), different types of immune cells (CD14^+^-cells, i.e., monocytes/macrophages ‘M’, and PBMCs activated towards T-cells ‘T’)] used in experiments for this publication are human primary cells which were isolated from subcutaneous skin biopsies and donor-specific blood samples. In case of co- and multi-cultures, experiments were always conducted in an autologous manner. Table 2 provides an overview of patient demographics and relevant medical records. Weight statuses were ranked according to World Health Organization (World Health Organization, n.d.).

Human subcutaneous skin and adipose tissue biopsies were obtained from plastic surgeries performed by Dr. Wiebke Eisler (BG Klinik Tübingen, Tübingen, Germany) and Dr. Ulrich E. Ziegler (Klinik Charlottenhaus, Stuttgart, Germany), approved by the local medical ethics committee: Patients gave an informed consent according to the permission of the “Landesärztekammer Baden-Württemberg” (No. 495/2018BO2 and F-2020-166). All procedures were carried out in accordance with the rules for medical research of human subjects, as defined in the Declaration of Helsinki. Collection of human whole blood samples was performed in accordance with the Declaration of Helsinki. Healthy blood donors and patients gave informed consent as approved by the Ethical Committee of the Eberhard Karls University Tübingen (No. 495/2018-BO02 for the isolation of PBMCs from whole blood).

Throughout the isolation, injection and chip culture processes, different cell-type specific cell culture media were used. All media, except for the endothelial cell medium, are defined compositions. Table 3 provides an overview of the different types of cell culture media used for cell-type specific isolations, pre-chip and on-chip cultures.

**Table 3.**
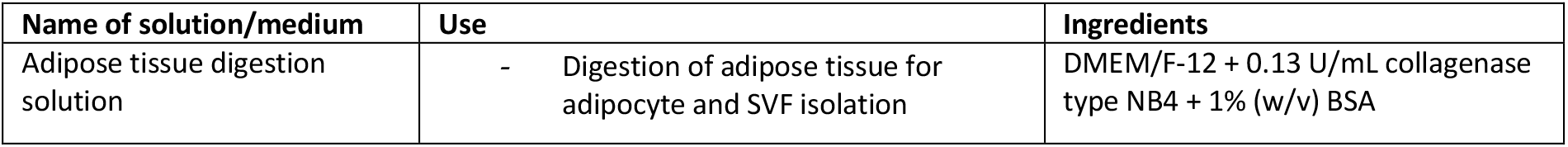
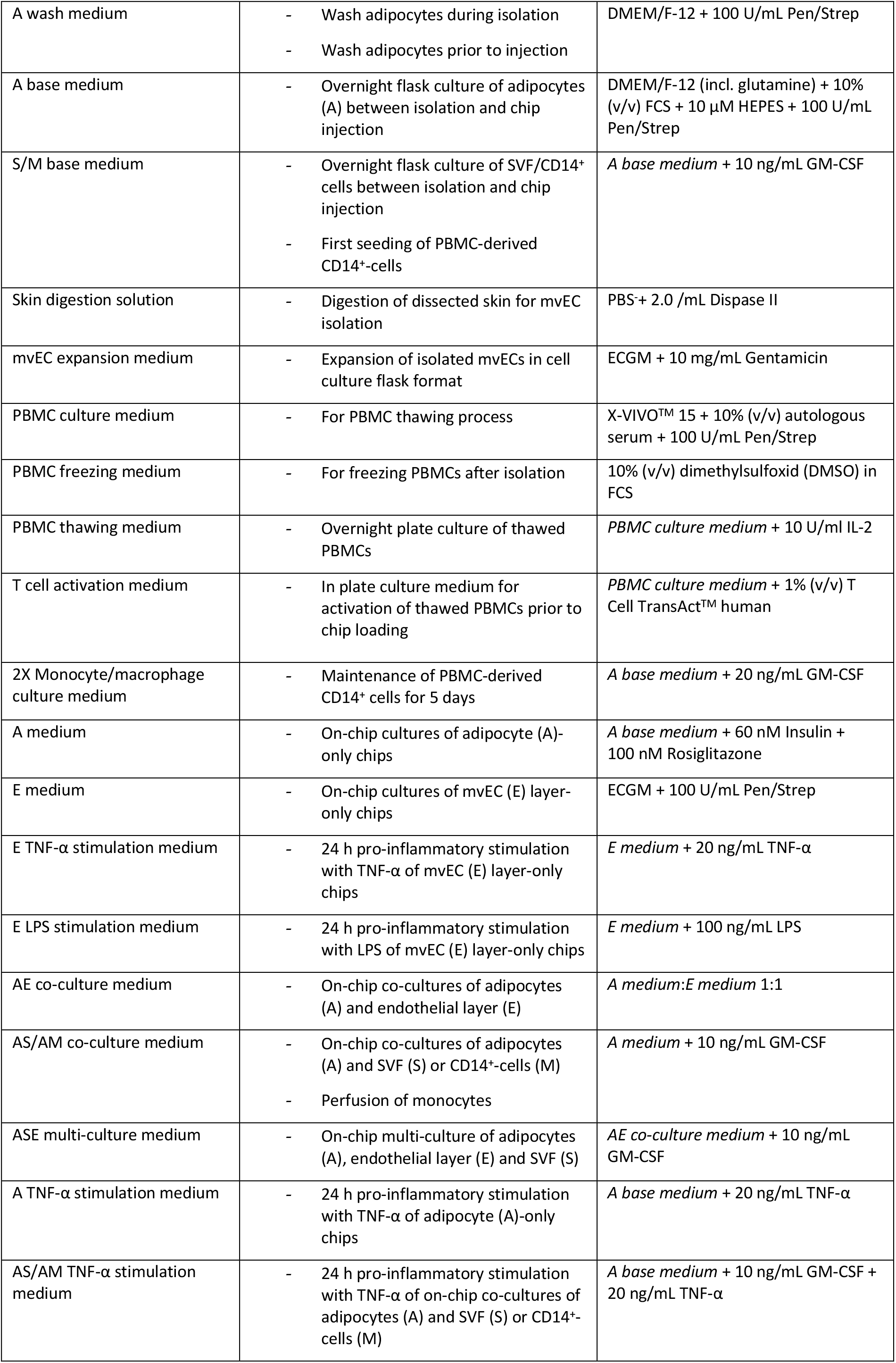
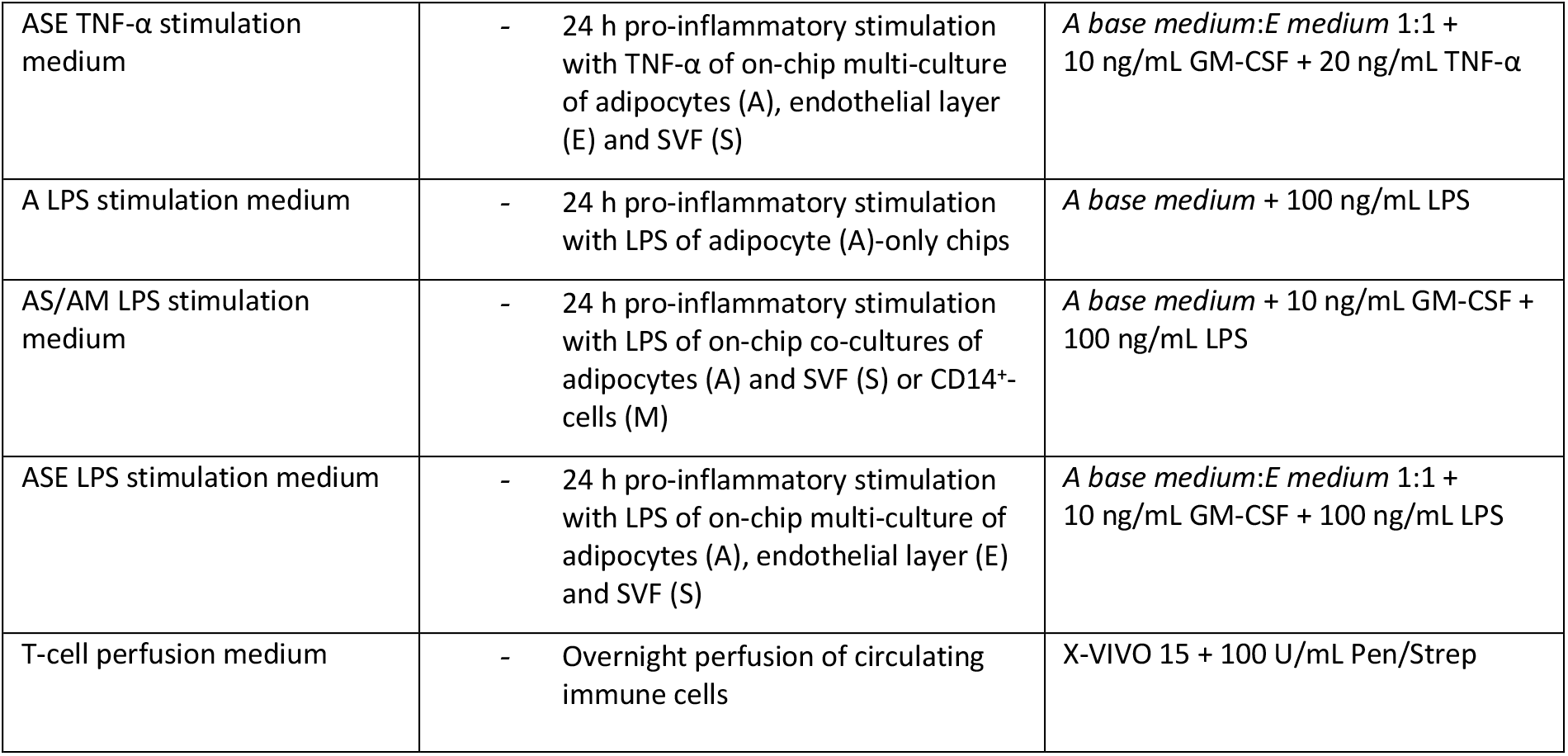
Overview of different types of enzymatic digestion solutions and cell culture media used in this study.

#### Isolation and pre-chip culture of mature adipocytes

Primary mature adipocytes were isolated from human skin and subcutaneous adipose tissue biopsies on the same day of surgery. We recently described our isolation process at length in a methodical book chapter (Rogal et al., 2021). In brief, the skin was separated from the subcutaneous adipose tissue and used for isolation of endothelial cells. The adipose tissue was then rinsed with Dulbecco’s phosphate buffered saline with MgCl_2_ and CaCl_2_ (PBS^+^; Merck KGaA, Darmstadt, Germany) twice, and macroscopically visible blood vessels and connective tissue structures were carefully removed. The remaining adipose tissue was cut into small pieces of approximately 1 cm^3^ and subsequently enzymatically digested by incubation in a collagenase solution (*adipose tissue digestion solution*) for 60 min at 37°C on a rocking shaker (50 cycles/min; Polymax 1040, Heidolph Instruments GmbH & CO. KG, Schwabach, Germany). Finally, the digested adipose tissue was strained through a mesh size of 500 µm and washed three times with DMEM/F-12, no phenol red (21041025; Thermo Fisher Scientific Inc., Waltham, MA) with 100 U/mL Penicillin/Streptomycin (Pen/Strep) (*A wash medium)*. For each washing step, adipocytes and medium were gently mixed, and left to rest for 10 min. After separation of the buoyant adipocytes and the medium, the liquid medium from underneath the packed layer of adipocytes was aspirated. Adipocyte isolation was performed on the day before injection into the chips. The freshly isolated adipocytes were cultured overnight, by adding equal volumes of packed adipocytes and *A base medium* to a culture flask kept in a humidified incubator at 37°C and a 5% CO_2_ atmosphere.

#### Isolation and pre-chip culture of mvECs

Human microvascular endothelial cells (mvECs) were isolated from resected skin from plastic surgeries. A piece of approximately 8 cm^2^ was washed and submersed in Phosphate Buffered Saline without calcium chloride and magnesium chloride (PBS^-^; L0615; Biowest, Nuaillé, France). Subcutaneous fat as well as big blood vessels were removed, and the remaining skin was cut in strips of approximately 4 cm length and 1 mm width and finally incubated in 10 ml *skin digestion solution* (2.0 U/mL dispase D4693, Merck KGaA, in PBS^-^) at 4°C overnight. The following day, the epidermis was peeled off using tweezers and the remaining strips of dermis were washed twice in PBS^-^. After a short incubation in Versene Solution (15040066; Thermo Fisher Scientific Inc.), dermis strips were incubated for 40 min in 0.05% Trypsin in EDTA Solution (59418C; SAFC) at 37 °C [trypsin reaction stopped by adding 10% Fetal Calf Serum (10326762; HyClone^TM^, Cytiva Europe GmbH, Freiburg, Germany)] to loosen the cells from the tissue. The strips were transferred to the lid of a petri dish containing 10 mL pre-warmed PBS^-^. Processing each dermis strip at a time, the dissociated cells were scraped out with a scalpel. After each strip was scraped for at least 8 times, the resulting cell suspension was strained (mesh size 70 µm) into a centrifuge tube and the petri dish lid was rinsed two more times with PBS^-^. To obtain a cell pellet, the cell solution in the centrifuge tube was centrifuged at 209 rcf for 5 minutes. The supernatant was discarded, and the pellet was resuspended in pre-warmed 10 mL Endothelial Cell Growth Medium (ECGM; C-22010, PromoCell GmbH, Heidelberg, Germany) with 10 mg/mL Gentamicin (*mvEC expansion medium*), seeded into two T25 cell culture flasks and incubated at 37°C, 5% CO_2_ and 95% rH overnight. On the next day, dead cells and debris were washed off by rinsing with PBS^-^ followed by addition of fresh *mvEC expansion medium*.

To remove fibroblasts from the expansion flask, the cells were washed with PBS^-^ and incubated in Versene Solution at 37°C, 5% CO_2_ and 95% rH until the fibroblasts detached. Following the aspiration of the Versene Solution, the cells were washed once again with PBS^-^ and pre-warmed *mvEC expansion medium* was added. Versene treatment was repeated accordingly throughout the first days of culture when needed. Else, media was changed every 3 days until the cells were injected into the microfluidic platform.

To (i) achieve sufficient cell count for chip injection and to (ii) purify isolated mvECs from fibroblast contamination as described above, mvECs must be expanded in flask format for at least 6 days after isolation. For experiments on endothelial layer-only chips (no donor-specificity required), mvECs from one donor were cryopreserved and re-used for the whole series of the experiment.

#### Isolation and pre-chip culture of SVF

Stromal vascular fraction (SVF) was isolated from human subcutaneous adipose tissue biopsies on the same day of surgery. The adipose tissue sample was rinsed with PBS^+^ twice, large blood vessels were carefully removed and then cut into small pieces of approximately 1 cm^3^. The adipose tissue pieces were then enzymatically digested by incubation in *adipose tissue digestion solution* (in equal volumes of adipose tissue and digestion solution) for 30 min at 37°C on a rocking shaker. After digestion, the tissue was passed through a strainer (mesh size: 500 µm) and left to rest for 10 min to allow for a separation of buoyant mature adipocytes, medium and non-buoyant cells. The packed layer of adipocytes was carefully aspirated, and the remaining cell suspension centrifuged for 5 min at 350 rcf. To lyse erythrocytes, supernatant was carefully decanted, and the cell pellet was gently re-suspended in Red Blood Cell Lysis Solution (freshly prepared according to manufacturer’s instruction; 130-094-183; Miltenyi Biotec B.V. & Co. KG, Bergisch Gladbach, Germany) which was incubated for 3 min at room temperature. Then, the cell suspension was strained through a 100 µm mesh size, collecting the filtrate in a centrifuge tube, and centrifuged for 5 min at 350 rcf. After decanting the supernatant, the cell pellet was resuspended in *S/M base medium* and cells were counted using Trypan blue and a hemocytometer. Cells from the SVF were cultured overnight in flask format in *S/M base medium* (seeding density of ∼1×10^5^ cells/cm^2^) or directly sorted *via* MACS to isolate CD14^+^-cells.

#### Peripheral blood mononuclear cell (PBMC) isolation, freezing and autologous serum collection

Isolation of fresh human PBMCs was initiated within 1 h after blood collection using Histopaque^®^ 1077 (10771; Merck KGaA) and standard density centrifugation (800 rcf, 20 min, no brakes). After centrifugation, PBMCs were washed twice in PBS^-^ supplemented with 0.1% BSA and 2 mM EDTA. PBMCs were used directly for isolation of CD14^+^-cells or immediately frozen at 10×10^6^ cells/mL in *PBMC freezing medium* using a CoolCell® Container (Corning).

For collection of autologous serum, whole blood was collected in S-Monovettes® containing serum gel with clotting activator (Sarstedt) followed by serum separation through centrifugation.

#### Isolation and pre-chip culture of CD14^+^-cells from SVF or PBMCs

CD14 is a co-receptor to the LPS receptor (lacking a cytoplasmatic domain, antibody binding, such as the MACS antibody, to CD14 alone does not provoke signal transduction) and is strongly expressed on monocytes and macrophages. To maintain cell identity and promote a monocyte-to-macrophage differentiation, we supplemented the cell culture medium with granulocyte-macrophage colony-stimulating factor (GM-CSF) for the entire culture period. CD14^+^-cells were isolated by magnetic activated cell sorting (MACS) with positive selection using CD14 MicroBeads (130-050-201; Miltenyi Biotec B.V. & Co. KG) from freshly isolated SVF or PBMCs according to supplier’s instructions. In brief, MACS buffer was prepared freshly before each isolation by diluting MACS BSA Stock Solution (130-091-376; Miltenyi Biotec B.V. & Co. KG) 1:20 in autoMACS Rinsing Solution (130-091-222; Miltenyi Biotec B.V. & Co. KG). For degassing, the MACS buffer was sonicated for 10 min. Counted, freshly isolated cell suspension from SVF (cf. Isolation and pre-chip culture of SVF) or from PBMCs was centrifuged for 10 min at 350 rcf at 4°C to avoid activation of monocytes. Then, the supernatant was aspirated completely, and the cell pellet was resuspended in 80 µl of MACS buffer and 20 µl of CD14 MicroBeads per ≤1×10^7^ total cells. After incubating for 15 min at 4°C, the cells were washed by adding 2 ml of MACS buffer per ≤1×10^7^ total cells and centrifuged for 10 min at 350 rcf at 4°C. In the meantime, an LS column (130-042-401; Miltenyi Biotec B.V. & Co. KG) was placed into the magnetic field of an QuadroMACS Separator (130-090-976; Miltenyi Biotec B.V. & Co. KG) and primed by rinsing with 3 ml of MACS buffer. Flow-through was collected in a 15 ml centrifuge tube underneath the column. For separation, the cell pellet was resuspended in 500 µl of MACS buffer per ≤1×10^8^ total cells and applied onto the column. Unlabeled cells were collected by subsequently washing the column by adding 3x 3 ml of MACS buffer. Then, the column was removed from the magnetic field and placed onto a new collection tube, 5 ml of MACS buffer were added onto the column, and magnetically labeled cells were flushed out by firmly pushing the plunger into the column. The cell suspension was centrifuged for 10 min at 350 rcf at 4°C. SVF-derived CD14^+^-cells were resuspended in *S/M base medium*, cells were counted and seeded at a density of ∼1×10^6^ cells/cm^2^ for overnight culture. PBMC-derived CD14^+^-cells were resuspended in *S/M base medium* and cultured at a density of ∼3×10^5^ cells/cm^2^. Every two days, 50 % of medium was exchanged with *2X Monocyte/macrophage culture medium*.

#### Thawing of PBMCs and activation of T-cells prior to chip culture

To thaw frozen PBMCs, cryopreserved cells were shortly placed at 37 °C, resuspended in prewarmed *PBMC culture medium* (X-VIVO^TM^ 15 medium (BE02-060F; Lonza Group AG, Basel, Switzerland) supplemented with 10% autologous serum and 100U/mL Pen/Strep), centrifuged and cultured at ∼1.5×10^6^ cells/cm^2^ in *PBMC thawing medium* overnight. PBMCs were washed and cultured in *PBMC culture medium* at a density of 0.5 – 1×10^6^ cells/mL in a total volume of 1 ml. CD3/CD28-mediated activation of T cells was conducted by using 10 μl of T cell TransAct^TM^ (130-111-160; Miltenyi Biotec B.V. & Co. KG) according to manufacturer’s instructions (*T cell activation medium*). T cells were activated for 3 days prior to injection.

### On-chip culture of adipose tissue

#### General remarks on injection, handling and readouts

Figure 7 provides an overview of the general timeline of WAT-on-chip experiments. Adipocyte-only as well as adipocyte-SVF and adipocyte-CD14^+^-cell co-culture chips were injected on the day after isolation. Day of chip injection is defined as day 0 (d0) for all experiments. After injection, on-chip tissues were supplied with respective culture media (cf. table 3) via gravity-driven flow overnight. On d1, chips were then connected to constant media perfusion via an external syringe pumping system (LA-190, Landgraf Laborsysteme HLL GmbH, Langenhagen, Germany). For connecting the chips to the syringe pump, we used Tygon tubing (0.762 x 2.286 mm, e.g. Tygon® ND 100-80 Medical Tubing, Saint-Gobain Performance Plastics Pampus GmbH, Willich, Germany), 21 GA stainless steel plastic hub dispensing needles (e.g., KDS2112P, Weller Tools GmbH, Besigheim, Germany; connected to Luer Lok™ style syringes) and blunt 21 GA stainless steel needles (made from the dispensing needles by removing the plastic hub after dissolving the glue overnight in a 70% ethanol solution). Media perfusion was realized in push mode, flow rate set to 20 µL/h. Unless stated otherwise, medium was changed every other day by re-filling inlet tubing and syringe reservoirs with fresh, pre-warmed culture medium. Endpoint analyses were conducted on d5 (d6, respectively, for monocyte perfusion) or d12 (d13, respectively, for T cell perfusion). In the case of stimulation experiments, effluents were collected for the 24 h-period prior to stimulation in order to assess basal secretion for each chip. Stimulation then occurred for 24 h from d4 to d5, or from d11 to d12, respectively.

**Figure 7.**
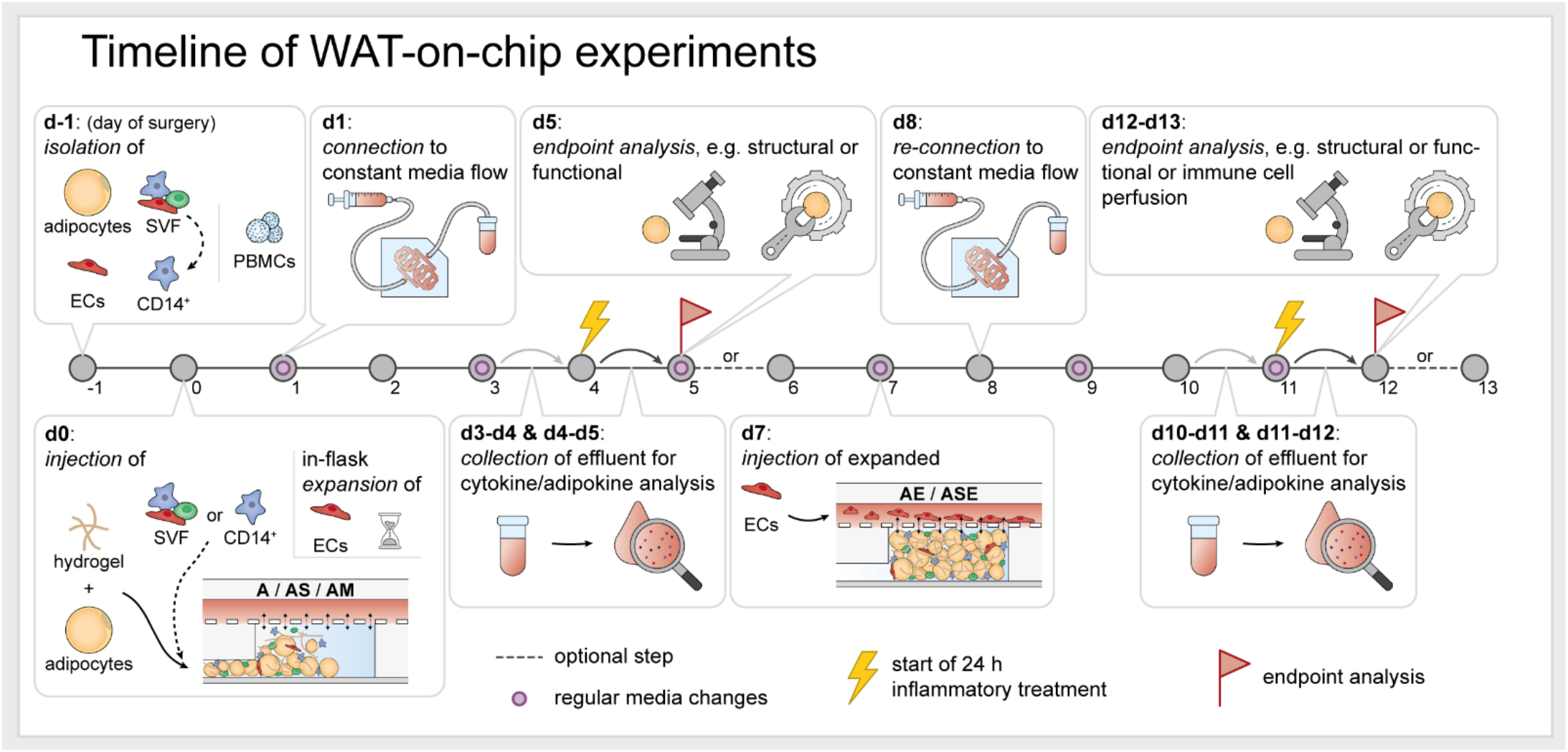
Overview of WAT-on-chip experiment timeline. On d-1, adipocytes, SVF and mvECs were isolated from skin/subcutaneous fat biopsies, and PBMCs were isolated from patients’ blood samples. D0 denotes the day of adipocyte (plus tissue chamber-resident cell types) injection. mvECs had to be expanded in culture flasks and could only be seeded onto the chips’ membranes on d7. After each injection, chips were supplied gravitationally overnight, before connecting to constant media perfusion on the next days. Media in pumping reservoirs were exchanged every other day. Most endpoint analyses were performed on d5 and d12.

#### Injection of adipose tissue into the microfluidic platform

Adipocytes were prepared by washing three times with *A wash medium* as described above (cf. Isolation and pre-chip culture of mature adipocytes). HyStem-C (GS313; CellSystems^®^, Troisdorf, Germany) hydrogel components were reconstituted and mixed according to manufacturer’s instructions (except for the crosslinker). Chips were prepared as described above (cf. Chip fabrication by soft lithography and replica molding) on the day before adipose tissue injection. Prior to injection, the chips were removed from PBS^-^ storage and a pipette tip filled with 100 µL PBS^-^ was inserted into the tissue compartment’s outlet to create a liquid droplet over the tissue inlet. Then, 60 µL of adipocytes from packed adipocytes layer were gently mixed with 25 µL of hydrogel components and 0.63 µL of Extralink (i.e., 40:1 hydrogel components:crosslinker). Then, 10 µL of the mixture were immediately injected into the chip’s tissue-chamber system by manual pressure. To avoid inclusion of air, the PBS-droplet over the tissue inlet port and adipocyte-hydrogel mixture were let coalesce before inserting the pipette tip into the port. Each tissue compartment system was loaded individually at a steady pace, to ensure that the adipocyte-hydrogel mixture reached the tissue chambers before the onset of gelation. When all tissue chambers were filled with adipocyte-hydrogel mixture, the injection channel was flushed with hydrogel by mixing 25 µL of hydrogel components with 6.25 µL of Extralink (i.e., 4:1 hydrogel components:crosslinker) and injecting 10 µL/chip via the tissue inlet ports into the tissue system. The injection ports were then closed using plugs. On-chip adipocytes were intermediately supplied by a gravitational media perfusion: an empty pipette tip was inserted into the media outlet port and a pipette tip filled with 100 µL of *A medium* was inserted into the media inlet port. Approximately 50 µL were manually pushed through the chip immediately to avoid crosslinking of the hydrogel inside the media channel. Using the method described above, up to 8 chips could be injected with one mixture before gelation of the hydrogel occurred. After overnight gravitational media supply, chips were connected to constant media perfusion of 20 µL/h. Media changes were performed every 3 days unless otherwise stated.

For injection of adipocyte-SVF or adipocyte-CD14^+^-cell co-culture chips, the above protocol was slightly adapted: adherent cells (i.e., SVF or CD14^+^-cells) were detached (see instructions below) and cell pellets of 0.5×10^6^ cells in 0.5 mL microcentrifuge tubes were prepared. Cell pellets were then resuspended in 25 µL of hydrogel components mix before adding 60 µL adipocytes and 0.63 µL of Extralink. Cell mixture injection, injection channel flushing, intermediate media supply and connection to constant media perfusion (using *AS/AM co-culture medium*) was done as described above for adipocyte-only chips.

Cells from the SVF were detached in sequential incubation steps with TrypLE™ Select Enzyme (1X) (12563011; Thermo Fisher Scientific Inc.): the growth area was rinsed once with PBS^-^. Then, the cell layer was incubated (37°C, 5% CO_2_ and 95% rH) for 5 min with 1:1 TrypLE™:PBS^-^, for 3 min with TrypLE™, and finally for another 8 min with TrypLE™. After each incubation step, the detachment solution was collected, and the enzymatic reaction was stopped by adding 10% (v/v) FCS to the cell suspension. Finally, the surface of the culture vessel was thoroughly rinsed to further detach cells. CD14^+^-cells were detached in a similar manner: 4 mg/mL lidocaine hydrochloride (L5647; Merck KGaA) were solved in Versene Solution freshly for each detachment. The growth surface was rinsed with PBS-once, and then the cells were detached by sequential incubation (37°C, 5% CO2 and 95% rH) in lidocaine solution for 3 min and 15 min. After each incubation step, the culture vessel was gently tapped from the bottom, and then the detachment solution was collected. After cell detachment, collected cell suspensions were pooled per cell type and centrifuged at 350 rcf for 5 min. After resuspension in *S/M base medium*, cells were counted, and transferred to microcentrifuge tubes as mentioned above.

#### Seeding of endothelial barriers in the microfluidic platform

Seven days after isolation, the mvECs were injected into the media channels of the microfluidic platforms to establish an endothelial barrier on the membrane separating tissue compartments from media perfusion.

For on-chip monoculture of endothelial layers, the chips’ tissue compartments were filled with HyStem-C (GS313; CellSystems^®^, Troisdorf, Germany) prior to mvEC seeding. Chips were prepared as described above (cf. Chip fabrication by soft lithography and replica molding) on the day before mvEC seeding. Prior to seeding, hydrogel components were reconstituted and mixed according to manufacturer’s instructions. The chips were removed from PBS^-^ storage and a pipette tip filled with 100 µL PBS-was inserted into the tissue compartment’s outlet to create a liquid droplet over the tissue inlet. Finally, 10 µL of hydrogel mixture were injected into the tissue compartment inlet port (to avoid enclosure of air during injection, liquid droplet over the inlet port and injection mixture were let coalesce before inserting the pipette tip into the PDMS). To avoid crosslinking of the hydrogel inside the media channel, 50 µL of PBS were flushed through the media compartment after hydrogel injection.

In case of multi-culture with other adipose tissue components, mvECs were added on d7 of adipocyte on-chip culture due to the required mvEC-expansion and -purification period described above. Immediately before EC injection into the co-culture systems, the media perfusion was disconnected by carefully removing the inlet and outlet tubing from the media ports of the system.

For mvEC detachment, medium was aspirated, and cells were washed with PBS^-^, and incubated with 0.05% Trypsin in EDTA solution (2 ml solution in T25 culture flask) for 5 min at 37°C, 5% CO_2_ and 95% rH. After 5 min, the enzymatic reaction was stopped by adding 10% FCS and the cell suspension was transferred to a centrifuge tube. The cell culture flask was rinsed once with PBS^-^. The cell suspension was centrifuged for 5 min at 209 rcf and the cell pellet resuspended in pre-warmed *mvEC expansion medium*. The cells were counted manually using a hemocytometer and the cell concentration was adjusted to 4×10^6^ cells/mL. A 100 µL filter tip was filled with 10 µL of the mvEC suspension and the tip was removed from the pipette. Carefully, the tip was inserted into the media inlet port of the chip. Introduction of air into the system was avoided by inserting the tip through a liquid droplet over the media inlet. An empty 100 µL filter tip was inserted into the media outlet and flow from the filled tip to the empty tip was ensured. The system was incubated for 2 h at 37°C to allow attachment of the mvECs. Within these 2 h, the mvEC suspension was gently moved inside the chip to increase membrane coverage by gently applying manual pressure on the pipette tips. After 2 h, the tips were removed carefully and 100 µL filter tips filled with 100 µL culture medium as defined by on-chip cell components (e.g., *E medium* for mvEC-only chips or *AE co-culture medium)* were inserted into media in- and outlet to provide static media supply at 37°C overnight. On the following day, the systems were (re-)connected to constant media perfusion. During the first 4 h, the media perfusion was ramped starting at 5 µL/h over the first 2 h, then 10 µL/h for 2 h, and finally set to 20 µL/h. Media changes were performed every 3 days unless otherwise stated.

#### Inflammatory stimulation

Inflammatory stimulations were performed by treating the chips for 24 h from d4-d5 or d11-d12 with TNF***-α*** (final concentration of 20 ng/mL; SRP3177; Merck KGaA) or LPS (final concentration of 100 ng/mL; 00-4976-93; Thermo Fisher Scientific Inc.) added to the respective media for each culture mode (table 3). To determine cytokine and metabolite concentrations in response to inflammatory stimulation, effluents were collected for the 24 h before treatment (baseline release for each chip) and after the 24 h-treatment.

#### Perfusion of immune cells

Activated T-cells were perfused for 18 h from d12-d13 of on-chip culture, and recruitment to different adipose tissue culture modes was studied (*A*, *AS*, *ASE*). T-cells were detached by (i) removing half of the culture medium, (ii) gently pipetting and collect already detached cells in a centrifuge tube, (iii) rinsing the growth surface with PBS^-^, gently pipetting and collecting the cell suspension again. Then, the cell suspension was centrifuged at 300 rcf for 5 min. Before perfusion, T-cells were labeled with CellTracker™ Deep Red Dye (C34565; Thermo Fisher Scientific Inc.) by resuspending the cell pellet in CellTracker Solution (reconstituted according to manufacturer’s instructions and further diluted to 2 µM in DMEM + 100 U/mL Pen/Strep) and incubating for 60 min (37°C, 5% CO_2_ and 95% rH). Labeled cells were then centrifuged at 300 rcf for 5 min, resuspended in X-VIVO^TM^ 15 + 100 U/mL Pen/Strep, counted and adjusted to a cell concentration of 375,000 cells/mL (à 275 µL/chip, i.e., circa 100,000 cells/chip). T-cells were perfused through the chips media channel by inserting a pipette tip containing 275 µL cell suspension into the media outlet and withdrawing the suspension with a flow rate of 10 µL/h. T cell recruitment to different adipose culture modes was quantified by determination of cell tracker fluorescence intensity in the tissue chambers.

Circulating monocytes derived from PBMCs were perfused for 24 h from d5-d6 of on-chip culture, and recruitment to one culture mode (*A*) was studied comparing 3 µm- and 5 µm membrane pore sizes. Before detachment, the CD14^+^-cells were labelled with CellTracker™ Deep Red Dye by incubating CellTracker Solution (reconstituted according to manufacturer’s instructions and further diluted to 2 µM in DMEM + 100 U/mL Pen/Strep) for 60 min (37°C, 5% CO_2_ and 95% rH). Afterwards, cells were washed by adding *S/M base medium*. Detachment was performed as described above (cf. Injection of adipose tissue into the microfluidic platform), and the cell concentration was adjusted to 112,500 cells/mL (250 µL/chip --> 28,125 cells/chip) in *AS/AM co-culture medium*. CD14^+^-cells were perfused through the chips media channel by inserting a pipette tip containing 250 µL cell suspension into the media outlet and withdrawing the suspension with a flow rate of 10 µL/h.

### Structural characterization of adipose tissue components on-chip

#### Endothelial barrier function assessment

Endothelial barrier integrity was assessed for the endothelial layer-only culture mode using a macromolecular tracer approach. For comparison, chips without endothelial barrier, only with a hydrogel gel-filled tissue compartment were measured. On d5 of on-chip culture, media supplemented with 100 µg/mL FITC-dextran with sizes of 3-5 kDa (FD4; Merck KGaA) or 40 kDa (FD40; Merck KGaA) were perfused through the chip at a flow rate of 20 µL/h for 60 min. Using a confocal Laser-Scanning-Microscope (LSM 710, Carl Zeiss Microscopy GmbH, Jena, Germany), fluorescence intensity was determined for 3 different focal planes (lower tissue chamber, upper tissue chamber and media channel) every 5 s. For analysis, we measured the mean grey value for each focal plane position for each time point using Fiji (Image J version 1.53c) (Schindelin et al., 2012), subtracted the background mean grey value and adjusted the offset in time it took for the tracer to reach the medium channel. Mean grey values were then normalized to the mean grey value measured at the final time point in the media channel.

#### Labelling for cell tracking

To trace and visualize the SVF, cells were labeled with a cell tracker prior to injection. Before detaching the cells, they were incubated in CellTracker™ Deep Red Dye solution (reconstituted according to manufacturer’s instructions and further diluted to 2 µM in DMEM + 100 U/mL Pen/Strep) for 60 min (37°C, 5% CO2 and 95% rH). Afterwards, cells were washed by replacing the labelling solution by *S/M base medium*.

#### (Immuno-) staining

A variety of (immuno-) staining procedures were performed on d5 or d6 (only for monocyte recruitment experiment), respectively, or d12 or d13 (only for T-cell recruitment experiment), respectively. We used conjugated and unconjugated antibodies.

All conjugated antibodies (table 4) were stained prior to fixation (except for CD11c and eNOS) by washing the chips once with PBS^+^ and twice with PBS^+^ with 0.5% (w/v) BSA. Then the antibody was diluted in PBS^+^ with 0.5% (w/v) BSA and 20 µM Hoechst 33342 Solution (62249; Thermo Fisher Scientific Inc.) and incubated for 30 min (37°C, 5% CO_2_ and 95% rH), followed by two washing steps with PBS^+^ with 0.5% (w/v) BSA. Afterwards, chips were imaged within 45 min (Leica DMi8 with incubation unit, Leica Microsystems) or fixed directly for further staining. CD11c and eNOS conjugated antibodies were added to secondary antibody solutions at concentrations listed in table 4.

**Table 4.**
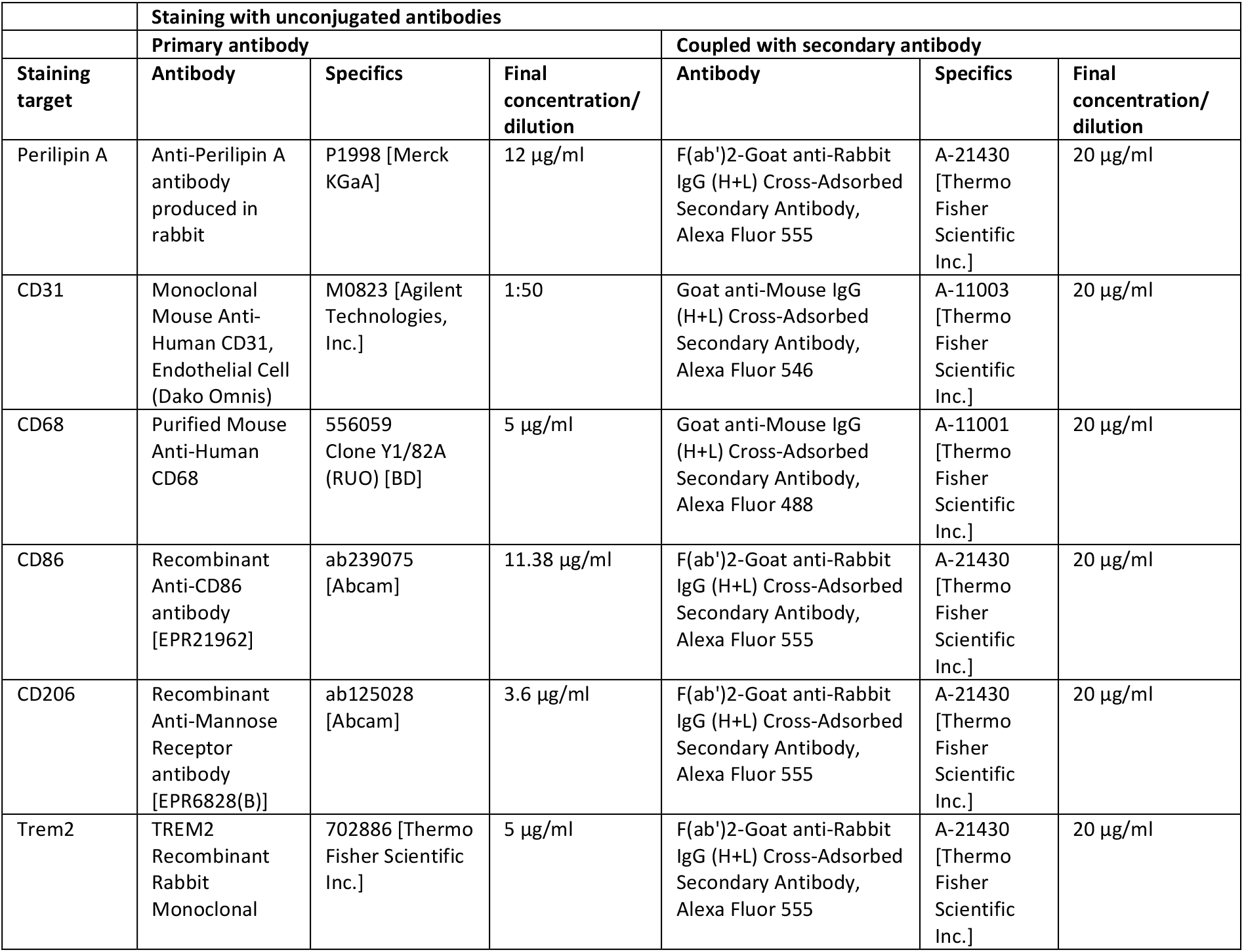
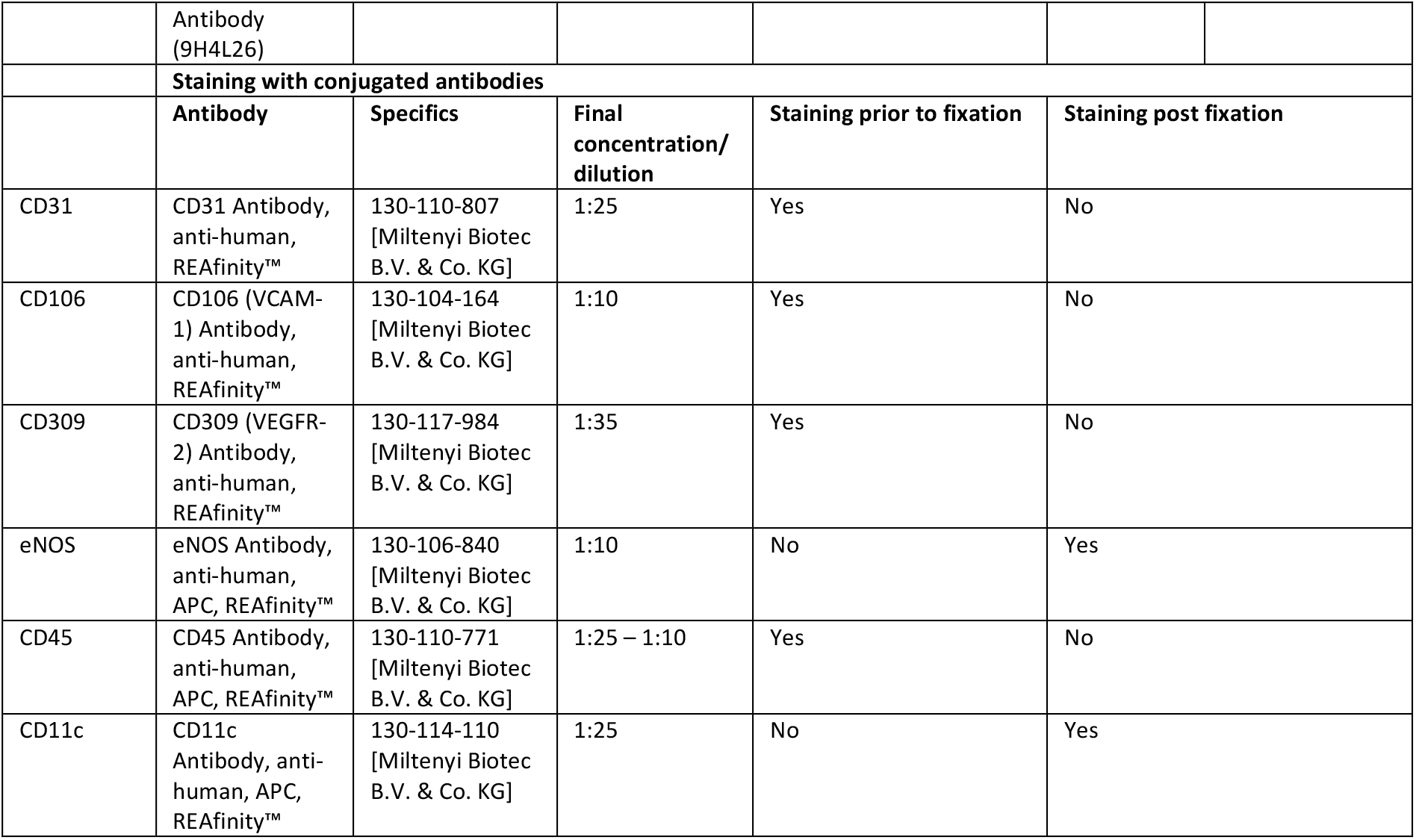
Overview of antibodies used for on-chip staining.

| Staining with unconjugated antibodies |  |  |  |  |  |  |
| --- | --- | --- | --- | --- | --- | --- |
| Primary antibody |  |  |  | Coupled with secondary antibody |  |  |
| Staining target | Antibody | Specifics | Final concentration/dilution | Antibody | Specifics | Final concentration/dilution |
| Perilipin A | Anti-Perilipin A antibody produced in rabbit | P1998 [Merck KGaA] | 12 µg/ml | F(ab')<sub>2</sub>-Goat anti-Rabbit IgG (H+L) Cross-Adsorbed Secondary Antibody, Alexa Fluor 555 | A-21430 [Thermo Fisher Scientific Inc.] | 20 µg/ml |
| CD31 | Monoclonal Mouse Anti-Human CD31, Endothelial Cell (Dako Omnis) | M0823 [Agilent Technologies, Inc.] | 1:50 | Goat anti-Mouse IgG (H+L) Cross-Adsorbed Secondary Antibody, Alexa Fluor 546 | A-11003 [Thermo Fisher Scientific Inc.] | 20 µg/ml |
| CD68 | Purified Mouse Anti-Human CD68 | 556059 Clone Y1/82A (RUO) [BD] | 5 µg/ml | Goat anti-Mouse IgG (H+L) Cross-Adsorbed Secondary Antibody, Alexa Fluor 488 | A-11001 [Thermo Fisher Scientific Inc.] | 20 µg/ml |
| CD86 | Recombinant Anti-CD86 antibody [EPR21962] | ab239075 [Abcam] | 11.38 µg/ml | F(ab')<sub>2</sub>-Goat anti-Rabbit IgG (H+L) Cross-Adsorbed Secondary Antibody, Alexa Fluor 555 | A-21430 [Thermo Fisher Scientific Inc.] | 20 µg/ml |
| CD206 | Recombinant Anti-Mannose Receptor antibody [EPR6828(B)] | ab125028 [Abcam] | 3.6 µg/ml | F(ab')<sub>2</sub>-Goat anti-Rabbit IgG (H+L) Cross-Adsorbed Secondary Antibody, Alexa Fluor 555 | A-21430 [Thermo Fisher Scientific Inc.] | 20 µg/ml |
| Trem2 | TREM2 Recombinant Rabbit Monoclonal | 702886 [Thermo Fisher Scientific Inc.] | 5 µg/ml | F(ab')<sub>2</sub>-Goat anti-Rabbit IgG (H+L) Cross-Adsorbed Secondary Antibody, Alexa Fluor 555 | A-21430 [Thermo Fisher Scientific Inc.] | 20 µg/ml |

|  |  |  |  |  |  |
| --- | --- | --- | --- | --- | --- |
|  | Antibody (9H4L26) |  |  |  |  |
| **Staining with conjugated antibodies** |  |  |  |  |  |
|  | **Antibody** | **Specifics** | **Final concentration/dilution** | **Staining prior to fixation** | **Staining post fixation** |
| CD31 | CD31 Antibody, anti-human, REAfinity™ | 130-110-807 [Miltenyi Biotec B.V. & Co. KG] | 1:25 | Yes | No |
| CD106 | CD106 (VCAM-1) Antibody, anti-human, REAfinity™ | 130-104-164 [Miltenyi Biotec B.V. & Co. KG] | 1:10 | Yes | No |
| CD309 | CD309 (VEGFR-2) Antibody, anti-human, REAfinity™ | 130-117-984 [Miltenyi Biotec B.V. & Co. KG] | 1:35 | Yes | No |
| eNOS | eNOS Antibody, anti-human, APC, REAfinity™ | 130-106-840 [Miltenyi Biotec B.V. & Co. KG] | 1:10 | No | Yes |
| CD45 | CD45 Antibody, anti-human, APC, REAfinity™ | 130-110-771 [Miltenyi Biotec B.V. & Co. KG] | 1:25 – 1:10 | Yes | No |
| CD11c | CD11c Antibody, anti-human, APC, REAfinity™ | 130-114-110 [Miltenyi Biotec B.V. & Co. KG] | 1:25 | No | Yes |

For all types of unconjugated staining, on-chip tissues were fixed, permeabilized and blocked prior to antibody incubation. In brief, the chips were washed by flushing the media channels three times with PBS^+^ before fixing the on-chip tissues with ROTI®Histofix 4% (P087.6; Carl Roth GmbH + Co. KG, Karlsruhe, Germany) for 60 min at room temperature (RT) under gentle rocking on a rocking shaker. After fixation, the chips were washed three times with PBS^-^, permeabilized with 0.2% (w/v) Saponin in PBS^+^ for 20 min at RT and blocked with 0.3% (v/v) Triton-X and 3% (w/v) BSA in PBS^-^ for 30 min at RT. All primary and secondary antibodies were diluted in antibody diluent (S3022; Agilent Technologies, Inc, Santa Clara, CA) to concentrations listed in table 4. Primary antibodies were incubated at RT overnight. Secondary antibody solutions were supplemented with 1 µg/mL DAPI solution (MBD0015; Merck KGaA) and 1 µg/mL BODIPY™ 493/503 dye (D3922; Thermo Fisher Scientific Inc.) and incubated for 1 h at RT. Finally, the chips were washed three times with PBS^-^ and imaged within the next 48 h using a confocal Laser-Scanning-Microscope (LSM 710, Carl Zeiss MicroImaging). Importantly, to confirm endothelial coverage over the tissue chambers, the chips had to be inverted for imaging, or else the adipocytes in the tissue chamber obscured the EC barrier.

### Functional characterization of adipose tissue on-chip

#### Live-/dead staining

To evaluate the viability of the mvECS forming the vascular barrier on chip, a live-/dead staining was performed and imaged via fluorescence microscopy. The cytosol of living cells was stained with fluorescein diacetate (FDA) (F7378; Merck KGaA); the nuclei of dead cells with propidium iodide (PI) (P4170; Merck KGaA). A stock solution of FDA was prepared by dissolving the powder in aceton (1 mg/mL). PI powder was dissolved in PBS^-^ (1 mg/mL). Stock solutions were stored protected from light and diluted right before the staining process. FDA and PI stocks were diluted in 838 µL PBS^+^, adding 27 µL of PI and 135 µL of FDA. To stain the mvEC barrier-on-chip, the chip was disconnected from tubing and the media channel was flushed with PBS^+^ via hydrostatic pressure created by inserting a filled pipette tip into the media inlet port and an empty tip into the media outlet port. After equilibration of the PBS^+^ level in the tips, they were replaced by a tip filled with 50 µL live-/dead-staining solution. After another incubation of 5 min at 37°C, 5% CO_2_ and 95% rH, the tips were removed, and the media channel was flushed two times with PBS^+^ as described above. Fluorescent imaging was conducted immediately after the staining using a Leica DMi8 (with incubation unit, Leica Microsystems, Wetzlar, Germany).

#### Fatty acid uptake monitoring of adipocytes

To assess fatty acid uptake properties of on-chip adipocytes, a medium-chain fatty acid (Dodecanoic Acid, C12; BODIPY™ 500/510 C1, C12; D3823, Thermo Fisher Scientific Inc.) or a long-chain fatty acid (Hexadecanoic Acid, C16; BODIPY™ FL C16, D3821, Thermo Fisher Scientific Inc.) were added at a concentration of 4 µM to the culture medium as defined by culture mode (table 3). The uptake of the fatty acids was monitored in real-time using a fluorescence microscope with incubation (Leica DMi8 with incubation unit, Leica Microsystems) for 60 min. Fluorescence images were acquired every 3 min for each position. To quantify fatty acid uptake, for each time point per position, we measured mean grey values in the tissue chamber and in the plain media channel as background using Fiji software.

#### Responsiveness to ß-adrenoreceptor agonists

ß-adrenergic stimulation was performed by adding (−)-Isoproterenol hydrochloride (I6504; Merck KGaA) to culture medium as defined by culture mode (table 3). Final concentration ranged from 1 µM to 100 µM. For each final concentration, a corresponding 1000X stock solution was prepared by dissolving the isoproterenol in PBS^-^. Isoproterenol responsiveness was read out after a 2 h feeding phase of on-chip adipocytes with the fluorescently labeled fatty acid (BODIPY™ 500/510 C1, C12; D3823, Thermo Fisher Scientific Inc.) by analyzing the release of fatty acids from the adipocytes. Moreover, glycerol secretion after 24 h of stimulation was determined (cf. Analyses of effluents).

#### Acetylated low density lipoprotein (AcLDL) uptake by endothelial layer

Low Density Lipoprotein from Human Plasma, acetylated and coupled to a DiI complex (DiI AcLDL; L3484; Thermo Fisher Scientific Inc.) was added to *E medium* or *E TNF-α stimulation medium* (following a 24 h-stimulation) at a final concentration of 1 µg/mL. Uptake solutions were pre-heated and administered to the chips via gravitational flow (empty pipette tip in media outlets, pipette tip filled with 50 µL uptake solution in media inlets) for 3 h at 37°C, 5% CO_2_ and 95% rH. Afterwards, nuclei were stained by adding Hoechst 33342 Solution (62249; Thermo Fisher Scientific Inc.) to the uptake solutions for 20 min at 37°C, 5% CO_2_ and 95% rH. Uptake solutions were removed from the chips by gravitationally washing with *E medium* or *E TNF-α stimulation medium*. Uptake was imaged within 45 min (Leica DMi8 with incubation unit, Leica Microsystems).

#### Analyses of effluents (cytotoxicity, glycerol release and cytokine secretion)

For all experiments involving analyses of effluents, we used chips without cells (but with hydrogel-filled tissue compartments) as negative controls. These chips were run in parallel to tissue-laden chips and handled identically. Media effluents were collected over 24 h periods.

After collection, effluents were centrifuged at 1942 rcf for 10 min. Supernatants after centrifugation were directly processed for viability assessment and afterwards stored at −80°C for up to 4 months. They were not thawed more than twice. Prior to performing assays, effluents from storage and all required assay reagents were brought to RT.

To quantitatively assess the on-chip tissues’ viability, we measured the release of lactate dehydrogenase (LDH) into the media effluents using the CytoTox 96^®^ Non-Radioactive Cytotoxicity Assay (G1780, Promega GmbH, Walldorf, Germany). The assay was performed in a 384-well plate according to the manufacturer’s instructions. To determine a Target Cell Maximum LDH Release Control, we lysed the on-chip tissues for the different culture conditions (*A*, *AS*, *ASE*; biological duplicates per condition) by incubating 1X Lysis Solution in the respective culture media for 2 h. The mean of the measured absorbance values was assumed to be the maximal LDH release possible for the given experimental set-up and set to 100%.

For quantitative enzymatic determination of glycerol secretion, we used Free Glycerol Reagent (F6428; Merck KGaA), and Glycerol Standard Solution (G7793; Merck KGaA) for a standard curve. In technical duplicates, 60 µL of effluent were mixed with 40 µL of Free Glycerol Reagent in a 96-well plate. After 10 min incubation at 37°C, 5% CO_2_ and 95% rH, absorption at 540 nm was measured using a plate reader (Infinite^®^ 200 PRO, Tecan Trading AG, Männedorf, Switzerland). For each assay run, a standard curve was generated to correlate absorbances to glycerol concentrations). Cytokines were determined by fluorescent bead-based multiplex sandwich immunoassays (LEGENDplex™ Human Angiogenesis Panel 1, 740697 and LEGENDplex™ Human Adipokine Panel, 740196; BioLegend, Inc., San Diego, CA) read by flow cytometry (Guava easyCyte 8HT, Merck KGaA) following the manufacturer’s manual. In brief, effluents were analyzed in technical duplicates and incubated with a cocktail of target-specific capture beads followed by an incubation with biotinylated detection antibodies and finally with streptavidin-phycoerythrin (SA-PE). For each assay run, a standard curve was generated to correlate fluorescence intensities to cytokine concentrations. Flow cytometry data were evaluated with the LEGENDplex Cloud-Based Data Analysis Software Suite (BioLegend). Gates were adjusted manually to find optimal differentiation between capture bead populations, and the same gating strategy applied to all assay runs.

### Image processing, data presentation and statistical analysis

Images were processed using Fiji (Image J version 1.53c) (Schindelin et al., 2012) to adjust brightness and contrast, create maximum intensity projections or orthogonal views of Z-stacks and to insert scale bars. For 3D rendering and stitching of tile scans, we used the ZEN software (ZEN 2.3 (blue edition), Carl Zeiss Microscopy GmbH).

All data is presented as mean ± SE if not stated otherwise with sample sizes (n) stated for each case individually. For quantifications feasible on chamber level, such as optical readout, n denotes number of chambers covered in analyses. For quantifications feasible on chip level only, such as all kinds of effluent analyses, for instance, n denotes number of chip replicates. Descriptive statistics and graphs were generated using OriginPro (Version 2021, OriginLab Corporation). For testing statistical significance, we performed unpaired *t* tests using the online *t* Test Calculator tool provided by GraphPad (https://www.graphpad.com/quickcalcs/ttest1/?format=50). P value and statistical significance are indicated for each case individually.

## Acknowledgements

The authors thank Dr. Ziegler (Klinik Charlottenhaus, Stuttgart) for the kind provision of human adipose tissue from plastic surgery, Cristhian Rojas, Dominic Baum, Lena Scheying and Verena Haug for their assistance with cell culture and chip fabrication.

The research was supported in part by the Fraunhofer-Gesellschaft internal programs Talenta Start (to J.R.) and Attract (601543, to P.L.), by the Federal Ministry of Education and Research (BMBF; 031L0247B to P.L.) as well as the European Union’s Horizon 2020 research and innovation program under the Marie Skłodowska-Curie grant agreements no. 812954 (to P.L.) and no. 845147 (to M.C.).

## Author contributions

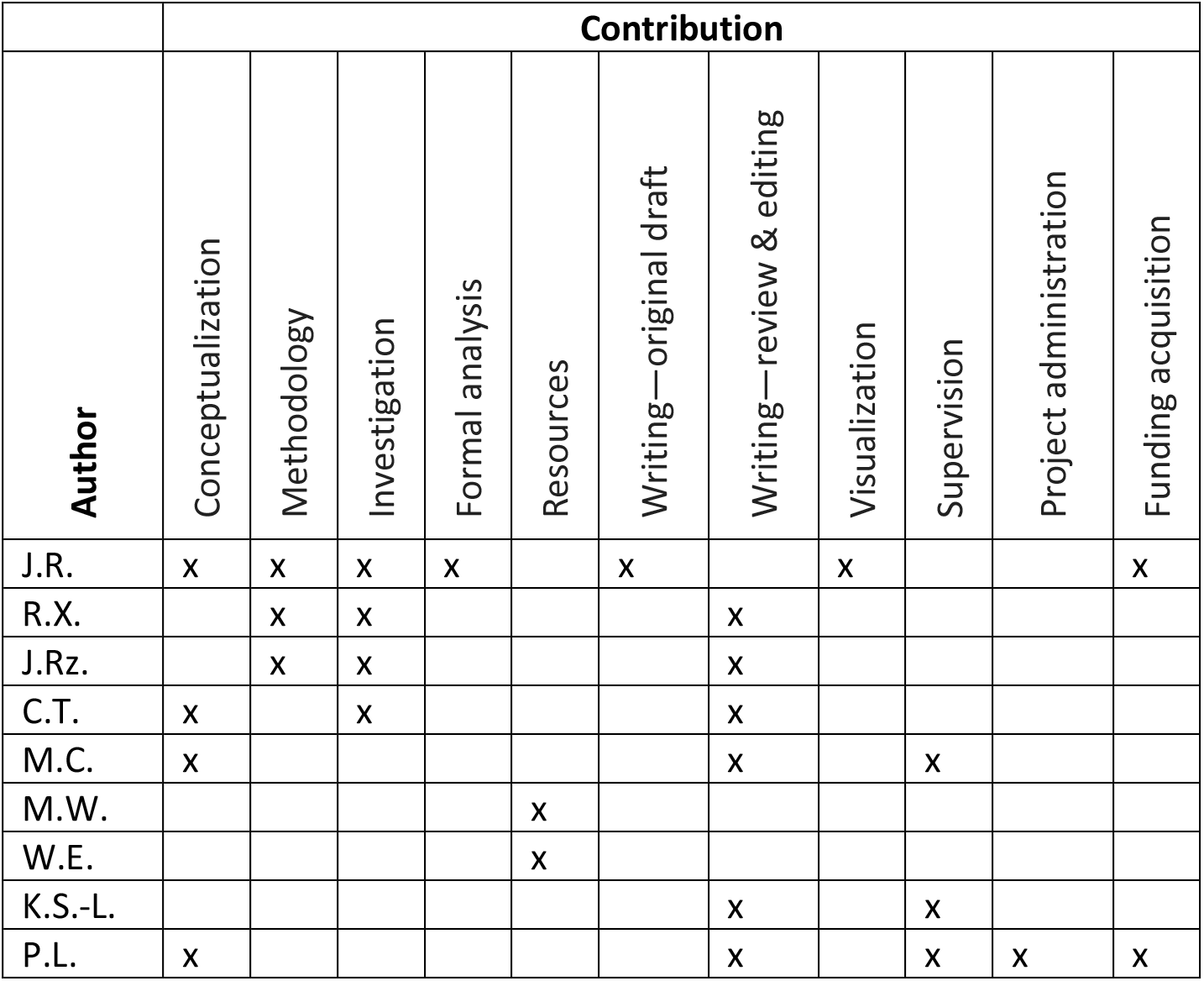

## Footnotes

1 Notably, conventional 3D volumetric imaging is particularly complicated for adipose tissue. Since obscuring effects from light scattering majorly occur at lipid-aqueous interfaces, imaging of adipocytes is highly susceptible to these effects, especially in deeper layers of the tissue.

