## supplementary figure for "Autologous human immunocompetent white adipose tissue-on-chip"

### Supplemental material

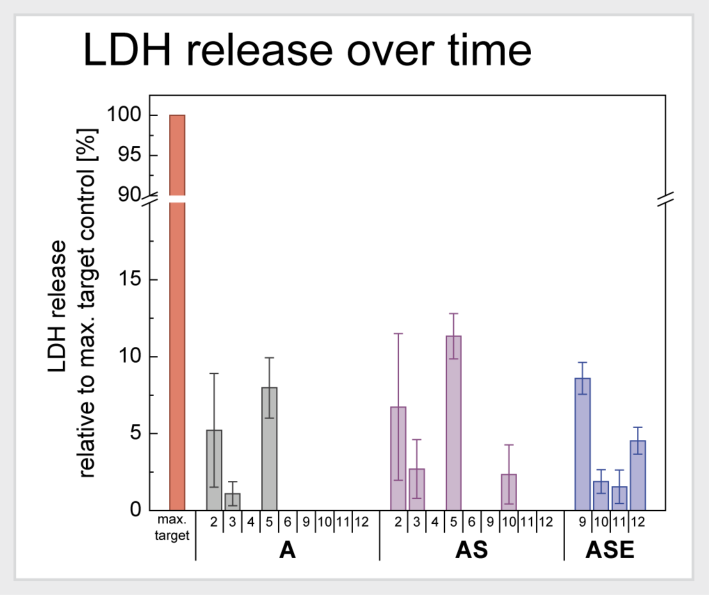


**Figure S1.** Cytotoxicity assessment for different culture modes during 12 d of on-chip culture. LDH release as cytotoxicity readout was determined every 24 h from media effluents. Absorbance values normalized to a respective target cell maximum LDH release control (100%), which was determined for each culture condition individually. On days that do not show bars, LDH release into the media effluents was not detectable with the readout method. Data were pooled from 3 donors. Due to different flow conditions, days 7 and 8 were omitted from the analysis. *A*: adipocyte-only chips; *AS*: adipocyte-SVF co-culture chips; *ASE*: adipocyte-SVF-mvEC co-culture chips.

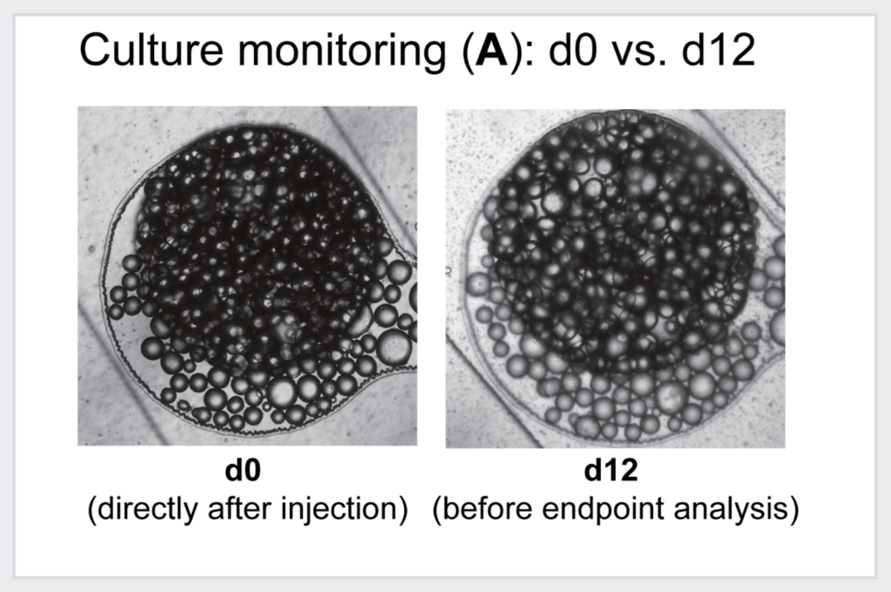


**Figure S2.** Monitoring of adipocytes-on-chip throughout a 12-day culture period. The same tissue chamber is shown directly after cell injection (d0) and directly prior to endpoint analysis (d12). Adipocyte morphology appeared comparable over time with slight deterioration regarding adipocyte stability towards d12.

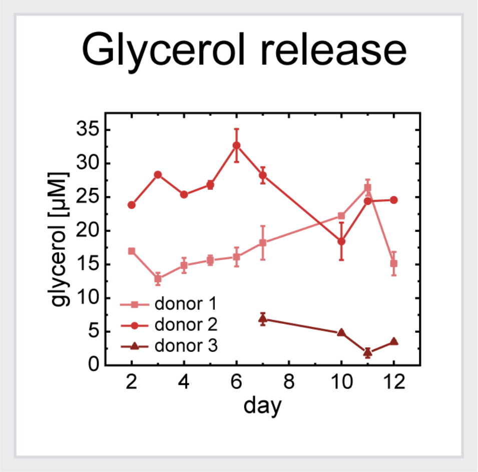


**Figure S3.** Monitoring of glycerol release from adipocytes into media effluents over time (donors 1 and 2 in biological duplicates; donor 3 in biological triplicates).

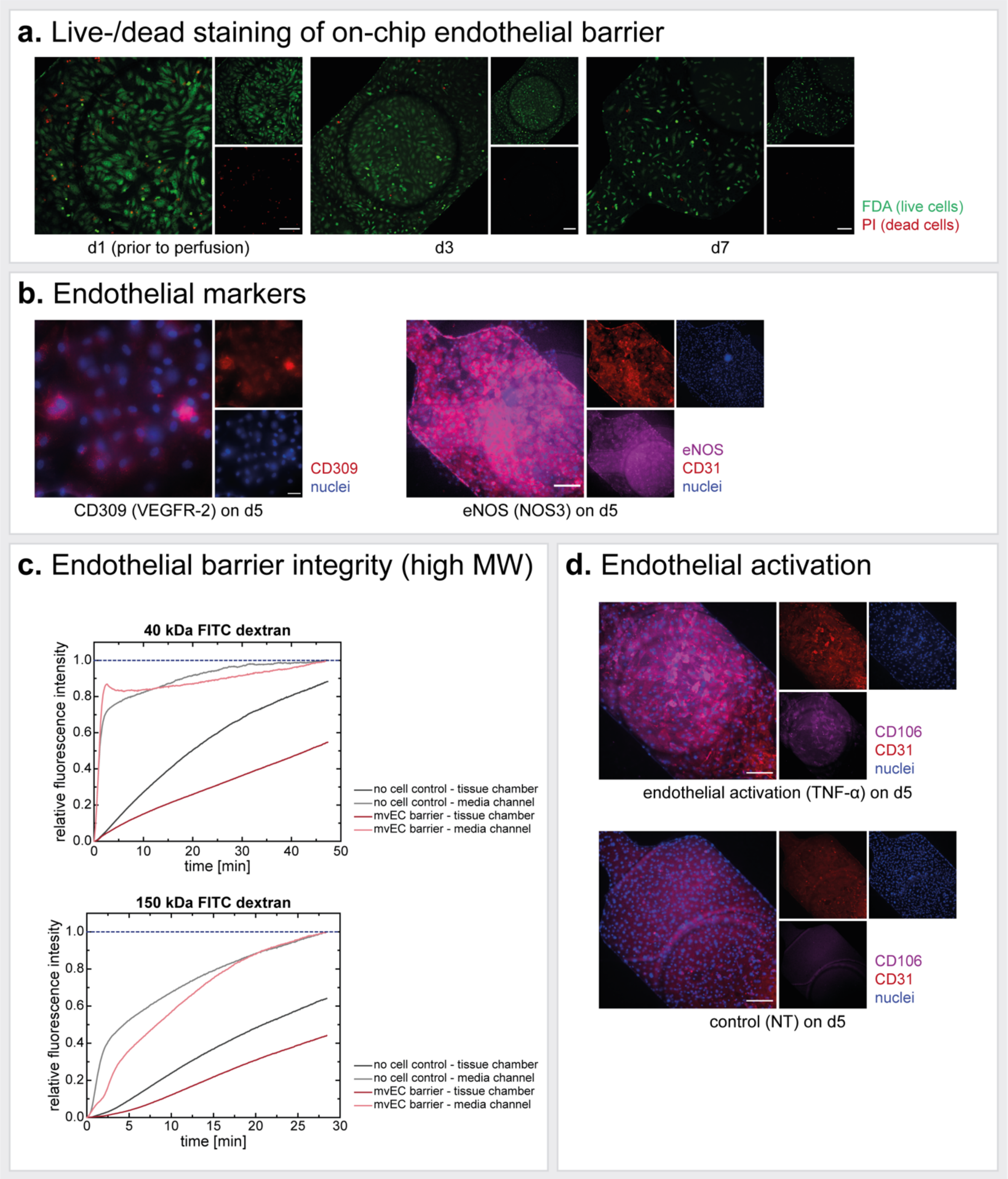


**Figure S4.** Additional characterization of on-chip endothelial barrier. (a) Live-/dead staining of on-chip mvEC layer on different days of analysis revealed an acceptable overall viability. On d1, prior to connection of constant media perfusion, there were several dead cells (potentially not fully attached remnants from the injection process). On d3 and d7, there were only a few dead cells in between the viable monolayer. Scale bars equal 200 µm. (b) In addition to CD31, we confirmed EC identity (and indicated proper functionality) by visualizing CD309 (alternatively VEGFR-2; main receptor of VEGF and important mediator of quiescent and active endothelium) (scale bar equals 50 µm) and eNOS (alternatively NOS3; for nitric oxide production) (scalebar equals 200 µm). (c) The permeability of the endothelial barrier on the chips' membranes was assessed by using fluorescent macromolecular tracers (here: 40 kDa and 150 kDa FITC-dextrans). The running time of the performed assays was not long enough to achieve equilibria between media channel and tissue chamber fluorescence intensity; yet there are clear trends indicating differences between cellularized vs. plain membrane transport as well as between different molecular weights. (d) Visualization of CD106 (alternatively VCAM1; expressed by activated endothelium for leukocyte-endothelial cell adhesion) for TNF- α-treated and untreated endothelial layers (scale bar equals 200 µm).

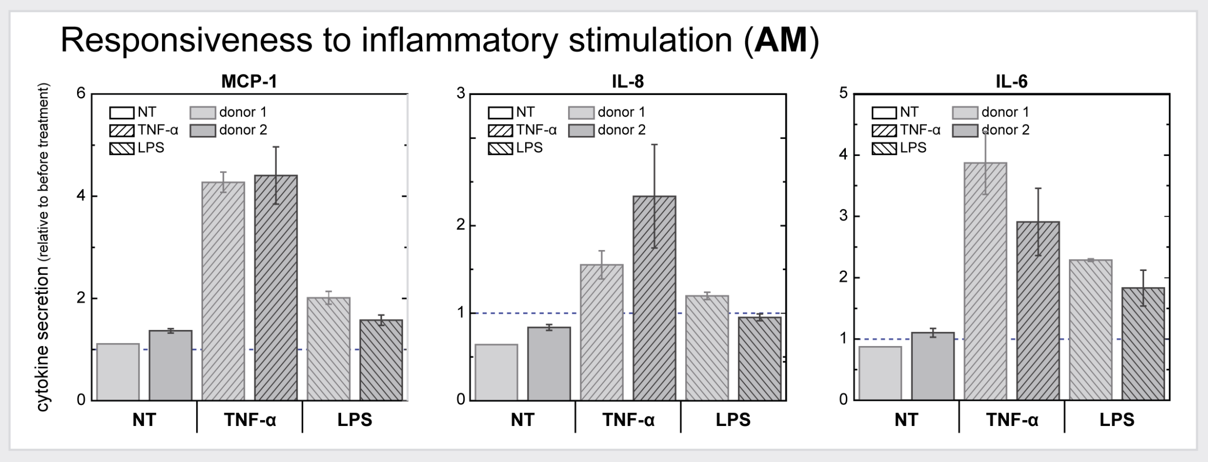


**Figure S5.** Proinflammatory cytokine release of adipocyte-CD14+-cell co-culture chips in response to TNF-$\alpha$ or LPS stimulation. Stimulation was performed for 24 h from d4-d5. Cytokine concentrations released throughout these 24 h were normalized to the cytokine levels determined for the 24 h before treatment for each chip. The experiment was performed for two different donors, and for two chips (i.e., biological duplicates) per condition.

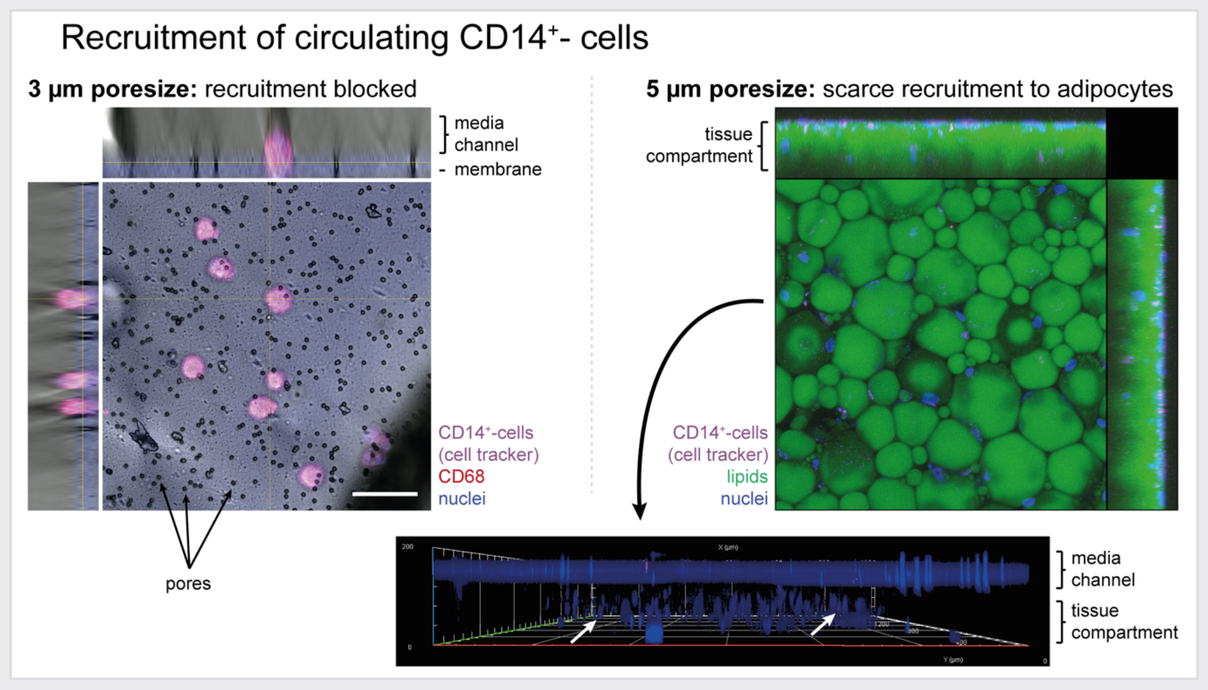


**Figure S6.** Comparison of recruitment of perfused CD14+-cells to adipocytes-on-chip (A) through 3 µm and 5 µm pore-sized membranes. CD14+-cells did not seem to be able to infiltrate the adipocyte chamber through 3 µm diameters pores in the chips' membranes. When building in 5 µm pore-sized membranes, a scarce recruitment into the tissue compartment could be detected as indicated by verification of cell tracker fluorescence signal in the tissue chamber.
